# CP204L Is a Multifunctional Protein of African Swine Fever Virus That Interacts with The VPS39 Subunit of HOPS Complex and Promotes Lysosome Clustering

**DOI:** 10.1101/2022.09.09.507285

**Authors:** Katarzyna Magdalena Dolata, Walter Fuchs, Grégory Caignard, Juliette Dupré, Katrin Pannhorst, Sandra Blome, Thomas C. Mettenleiter, Axel Karger

**Affiliations:** Institute of Molecular Virology and Cell Biology, Friedrich-Loeffler-Institute, Greifswald-Insel Riems, Germany; UMR Virologie, INRAE, Ecole Nationale Vétérinaire d’Alfort, Laboratoire de santé animale d’Alfort, Anses, Université Paris-Est, Maisons-Alfort, France; Institute of Diagnostic Virology, Friedrich-Loeffler-Institut, Greifswald-Insel Riems, Germany

**Keywords:** African swine fever virus, ASFV, CP204L, VPS39, HOPS complex, virus-host interaction, lysosomes

## Abstract

Virus replication depends on a complex interplay between viral and host proteins. In the case of African swine fever virus (ASFV), a large DNA virus, only few virus-host protein-protein interactions have been identified to date. In this study, we demonstrate that the ASFV protein CP204L directly interacts with the cellular homotypic fusion and protein sorting (HOPS) protein VPS39, blocking its association with the lysosomal HOPS complex, that modulates endolysosomal trafficking and promotes lysosome clustering. Instead, VPS39 is targeted to the sites of virus replication termed virus factories. Furthermore, we show that loss of VPS39 reduces the levels of virus proteins synthesized in the early phase of infection and delays ASFV replication but does not completely inhibit it. Collectively, these results identify a novel virus-host protein interaction that modulates host membrane rearrangement during infection and provide evidence that CP204L is a multifunctional protein engaged in distinct steps of the ASFV life cycle.

**Importance:** African swine fever virus (ASFV) was first identified over a hundred years ago. Since then, much effort has been made to understand the pathogenesis of ASFV. Yet, the specific roles of many individual ASFV proteins during the infection remain enigmatic. This study provides evidence that CP204L, one of the most abundant ASFV proteins, modulates endosomal trafficking during virus infection. Through direct protein-protein interaction, CP204L prevents the recruitment of VPS39 to the endosomal and lysosomal membranes, resulting in their accumulation. Consequently, CP204L and VPS39 become sequestered to the ASFV replication site. These results uncover a novel function of viral protein CP204L and extend our understanding of complex interaction between virus and host.

## Introduction

African swine fever virus (ASFV) causes a contagious and often lethal disease of domestic pigs and wild boars. The disease was first reported in Kenya in 1921 (1) and has remained endemic in Africa. Over the years, sporadic outbreaks were registered outside of Africa, but only in 2019 African swine fever (ASF) reached a pandemic level. There has been so far no evidence of cross-species transmission of ASFV to humans or mammals other than members of the family *Suidae*. Nevertheless, due to high case fatality rates approaching 100% in Eurasian suids and the lack of vaccines, the economic consequences of ASF are very high (2–4).

ASFV is a large double-stranded DNA virus of the family *Asfarviridae* that induces the synthesis of over 100 virus proteins in infected cells (5, 6). About 68 proteins are incorporated into virions (7). The virus replicates mainly in porcine monocytes and macrophages, although other cell types can be infected, especially in the later stages of the disease (8, 9).

ASFV utilizes multiple strategies to enter the host cell, including (i) binding to a hitherto unknown receptor followed by clathrin-mediated endocytosis (10), (ii) macropinocytosis (11), and (iii) phagocytosis (12). Once internalized, viral particles are trafficked along the endocytic pathway from peripheral early endosomes to late perinuclear endosomes (13). The acidic environment in late endosomes destabilizes the outer viral capsid and exposes its inner membrane, allowing the virus to fuse with the endosomal membrane and release the viral core with the genomic DNA into the cytoplasm (14, 15).

Similar to several other large DNA viruses, such as poxviruses and iridoviruses, the replication of ASFV is associated with cytoplasmic foci, referred to as virus factories (VFs). ASFV-induced VFs appear as complex and dynamic perinuclear structures close to the microtubule organizing center, surrounded by mitochondria (16) and a vimentin cage (17). ASFV factories are highly compartmentalized to coordinate different steps of the viral life cycle, such as virus DNA replication, transcription and translation, and virion assembly (16, 18). Importantly, VFs compartmentalization may protect the virus from degradation by antiviral defense mechanisms of the host cell. Despite their importance, the morphology of VFs and the mechanisms that lead to changes in the cellular organization required to produce complex replication sites are not yet understood.

It is known that ASFV reorganizes endosomal trafficking for its journey towards the perinuclear site, but endosomal membranes are also recruited to early VFs (19). The exact role of endosomal components in ASFV assembly is unknown. It has been suggested that endosomal compartments could be required for virus replication by providing a scaffold and confining the replication process to a specific cytoplasmic location. On the other hand, endosomal membranes may serve as intermediates for virus assembly. The virus-host interaction that leads to endosome accumulation in the early VF must occur in the initial phase of infection. Thus, characterizing interactions between host proteins and ASFV early proteins, synthesized before viral DNA replication, could shed light on the mechanism of VF assembly and transport of endosomal membranes into the factory.

The *CP204L* gene is conserved in all ASFV isolates and encodes a highly antigenic viral protein (20) essential for viral replication (21). The CP204L protein (hereinafter referred to as CP204L), also known as P30 or P32, is one of the most abundant viral proteins synthesized early during infection (22–24). In infected cells, CP204L is mainly localized in the cytoplasm, but small amounts have also been detected in the nucleus and at the plasma membrane (25). The nuclear CP204L has been reported to interact with the heterogeneous nuclear ribonucleoprotein K (hnRNP-K) (26). However, nothing is known about the interacting partners for the predominant cytoplasmic CP204L. Additionally, the molecular mechanisms by which CP204L interacts with the host and influences virus replication remain unexplored.

This study identifies a set of novel cellular and viral protein interactors of ASFV CP204L. In particular, we focus on host vesicular trafficking proteins, which are the key factors mediating ASFV infection progression. We demonstrate that CP204L directly interacts with VPS39, a component of the homotypic fusion and vacuole protein sorting (HOPS) complex, causing VPS39 dissociation from the HOPS complex and lysosome clustering. We discover that CP204L is recruited to VFs at the early times during infection. Moreover, our observations suggest that VPS39 is an important host factor that regulates the early steps of infection, but it is not essential for ASFV replication.

## Results

### Identification of the ASFV CP204L interactome

To gain insight into the host protein interactome of CP204L, we employed an affinity tag-purification mass spectrometry (AP-MS) approach (Fig. 1A). The CP204L of the highly virulent ASFV Georgia (27) with C-terminal GFP tag was used as a bait. The protein was stably expressed in wild boar lung (WSL) cells which were used for AP-MS experiments with and without ASFV infection. GFP-expressing WSL cells were used as a negative control. Proteins were affinity purified in biological triplicates (at minimum) and subjected to analysis by mass spectrometry to identify co-purifying partners. To minimize the false-positive identifications, host proteins bound to GFP in the absence of a bait protein CP204L were excluded from further analysis. Comparing the identified cellular interactors of the CP204L in infected and mock-infected cells revealed an overlap of 578 interactions (Fig. 1B). Additionally, 239 co-purified proteins were identified exclusively in virus-infected cells and 215 in cells without virus infection (see Table S1). Among the overlapping proteins, only five showed significant changes in protein levels after infection (Fig. 1C). To further functionally characterize the co-purifying proteins, we performed a Gene Ontology (GO) enrichment analysis via the traditional over-representation statistical method. Proteins interacting with the CP204L in mock and infected cells were enriched for several broad terms, such as cellular respiration, mitochondrial transport, and vesicle-mediated transport (Fig. 1D; Table S2). This latter category included a set of proteins involved in cellular processes critical for virus entry, immune evasion, and cell-to-cell spread, like endocytosis, autophagy, or retrograde trafficking. We therefore constructed a protein interaction subnetwork and looked specifically into interactions between CP204L and host proteins involved in vesicle transport, as well as interactions between CP204L and other ASFV proteins (Fig. 1E). From the host-virus interactions, one between the ASFV CP204L and the swine protein VPS39, a subunit of HOPS (homotypic fusion and vacuole protein sorting) complex, was notable in that it was identified with the highest abundance (log10 iBAQ; Table S3). The HOPS complex plays a role in endosomal cargo trafficking by mediating endosomal maturation (28) and fusion of lysosomes with late endosomes, phagosomes, or autophagosomes (29, 30). Of the protein-protein interactions (PPIs) among viral proteins, the A137R protein was the most enriched ASFV protein interacting with CP204L (Table S3 and Fig. S1). Unfortunately, we could not confirm the interaction between CP204L and A137R by an inverted pulldown (Fig. S2). Therefore, we focused on characterizing the interaction of CP204L with VPS39.

**Fig 1.**
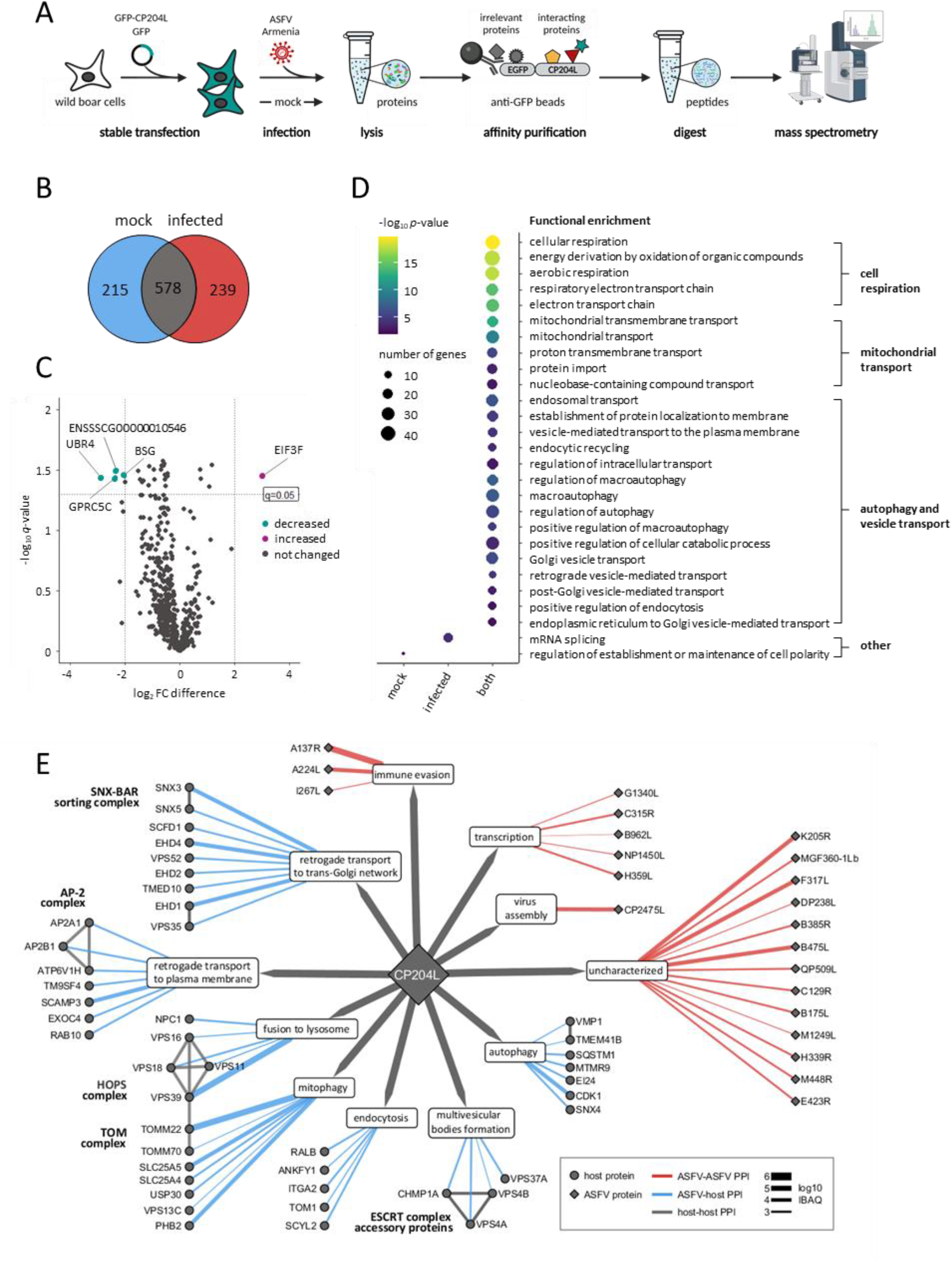
Identification of ASFV CP204L interactome. (A) AP-MS experimental workflow for identifying interactions between CP204L and host proteins. (B) Venn diagram demonstrating the overlap between protein interaction partners for CP204L identified in mock and virus-infected WSL cells. (C) Volcano plot showing differences in the abundance of overlapping host protein interactors. Dotted vertical and horizontal lines indicate the chosen cutoffs for fold-changes (FC, (|log2(FC)| > 2) and *q*-values (*q* < 0.05), respectively. Proteins showing significant differences in abundance between mock and virus-infected cells are marked by their gene names. (D) Selected functional GO terms from overrepresentation analysis are shown for each dataset. The most enriched terms are related to cell respiration, mitochondrial transport, autophagy and vesicle transport, and other terms. The color scale indicates significance expressed as -log10 *p*-value, and the size of the dots reflects the number of input genes associated with the respective GO term. (E) Network illustrating the interactions of CP204L with other ASFV proteins and host proteins involved in vesicle transport in the cell. Each node represents a protein (circles: host proteins; diamonds: ASFV proteins). Each edge is colored according to the type of interaction (blue: ASFV-host PPIs; red: ASFV-ASFV PPIs). Edge thickness is proportional to the log10 iBAQ value. Physical interactions among host proteins and their specific cellular functions were curated from the literature (Table S3). All shown CP204L-host interactions were identified in mock and virus-infected cells. CP204L-virus interactions were identified only in infected cells.

**Fig S1.**
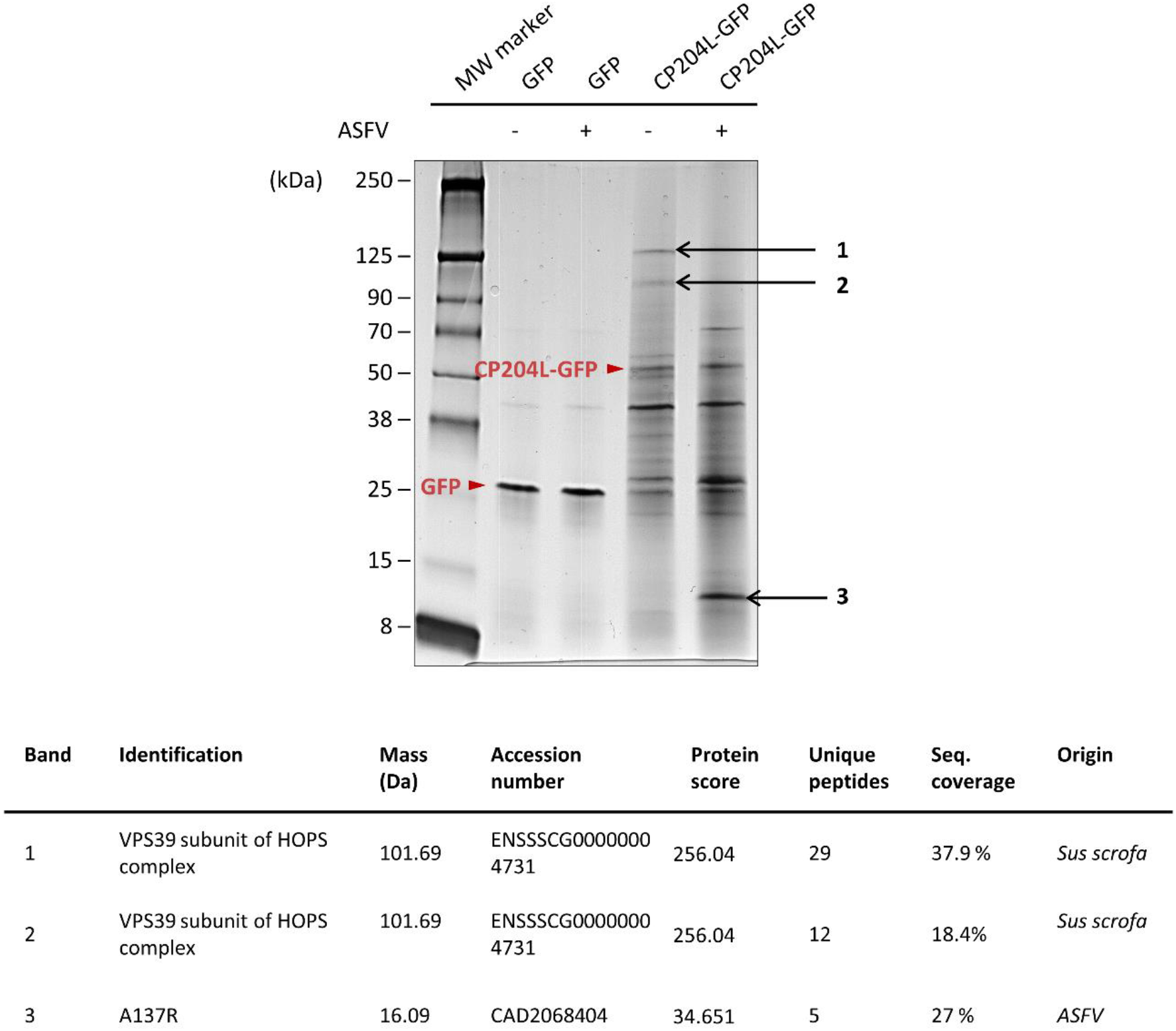
Silver stained SDS-PAGE of GFP affinity-purifications and protein identification by mass spectrometry. Eluates of GFP affinity purifications from mock and ASFV-infected WSL cells were subjected to SDS-PAGE and stained with silver. Bands 1-3 were excised from the gel and prepared for identification with mass spectrometry (Table S4; Text S1). The GFP and GFP-tagged CP204L are marked with red arrowheads.

**Fig S2.**
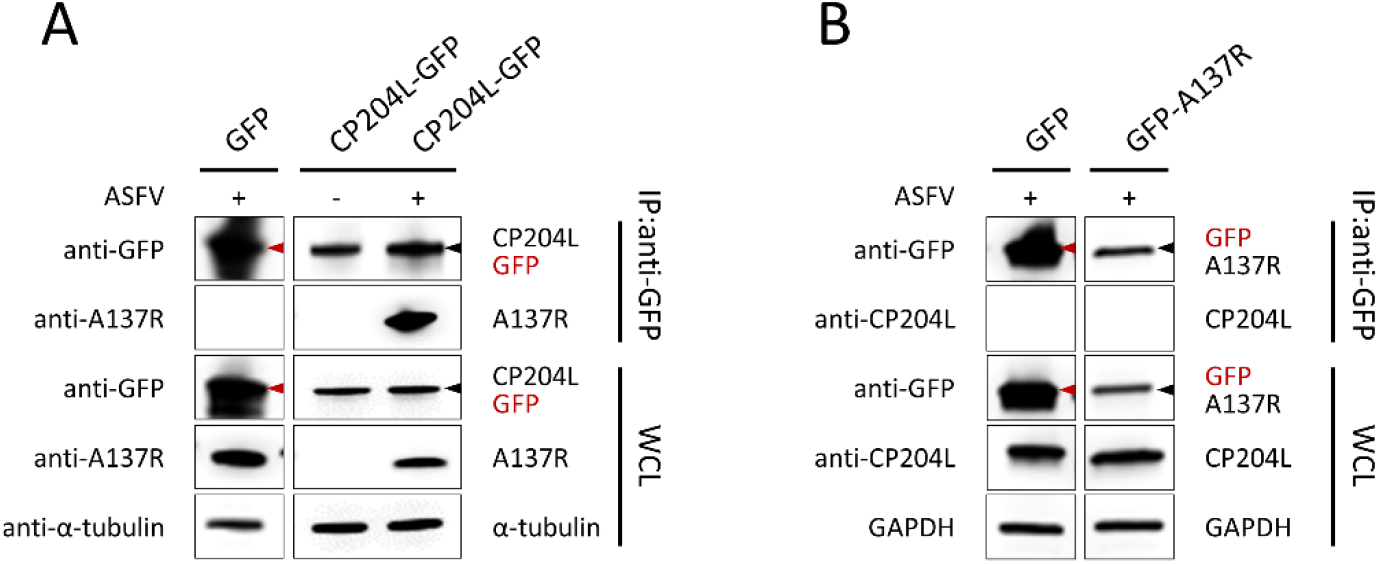
Reverse pulldown failed to confirm the interaction between ASFV CP204L and ASFV A137R. (A) Co-immunoprecipitation of CP204L-GFP with A137R in ASFV-infected cells. Lysates from mock and ASFV-infected cells stably expressing CP204L-GFP or GFP alone were subjected to GFP-specific immunoprecipitation. Representative immunoblots of whole cell lysates (WCLs) and GFP-immunoprecipitates (IP) are shown. α-tubulin was used as a loading control in WCLs. (B) Reverse co-immunoprecipitation of GFP-A137R with CP204L in ASFV-infected cells. No interaction between GFP-A137R and CP204L was detected (no band was present in the IP fraction). GFP was used as a control. GAPDH was used as a loading control in WCLs.

**Table S1**. All PPIs identified in mock- and virus-infected WSL cells expressing CP204L-GFP as bait.

**Table S2**. List of significantly (adjusted *p*-value < 0.01) enriched GO terms among identified host protein interactors of ASFV CP204L.

**Table S3**. List of selected CP204L interacting proteins: host proteins involved in vesicular transport and ASFV proteins.

**Table S4**. List of proteins identified from the silver-stained SDS-PAGE gel slices.

### CP204L interacts with the 241-541 amino acid region of the VPS39 subunit of HOPS complex

We performed a co-immunoprecipitation (co-IP) experiment with anti-GFP agarose beads in the presence of GFP-tagged CP204L or GFP alone as a control. Endogenous VPS39 co-precipitated with CP204L-GFP in uninfected and infected cells, whereas no interaction was observed between VPS39 and GFP (Fig. 2A). Reverse co-IP experiments with GFP-tagged VPS39 in infected cells further confirmed that CP204L and VPS39 interact specifically (Fig. 2B). To determine whether the CP204L-VPS39 interaction is direct, we applied a yeast two-hybrid (Y2H) assay. For this purpose, VPS39 and CP204L were fused to either the Gal4 DNA binding domain (DBD) or the Gal4 activation domain (AD), and fusion proteins were expressed in yeast as bait or prey, respectively. Physical interaction between CP204L-AD and VPS39-DBD proteins was confirmed by the Y2H assay (Fig. 2C). Moreover, both proteins showed the ability to form homodimers using the Y2H system. To further map the region of VPS39 required for the interaction with CP204L, gap repair cloning was applied. Forward and reverse primers were designed for every 270 nucleotides along the VPS39 sequence (corresponding to 90 amino acids). Fragments of VPS39 were introduced into the Gal4-DBD vector and co-expressed with CP204L fused to Gal4-AD in yeast cells grown on a selective medium. The results of this experiment indicated a 270-residue region encompassing position AA271 to AA541 of VPS39 protein to be critical for the interaction with CP204L (Fig. 2D). According to the InterPro (31) prediction, the CP204L binding region of VPS39 contains a clathrin heavy chain repeat (CHCR) domain. It also overlaps with a citron homology (CNH) domain located at the N-terminus and with a transforming growth factor beta receptor-associated domain 1 (Fig. 2E). Furthermore, an intrinsically disordered region is situated within the CP204L binding domain between AA440 and AA460 of VPS39, as predicted by IUPred2A (32). It is worth noting that we performed a similar domain mapping experiment for CP204L (Fig. S3). We could demonstrate that the middle region of CP204L is required for interaction with VPS39; however, the minimum interacting domain could not be specified.

**Fig 2.**
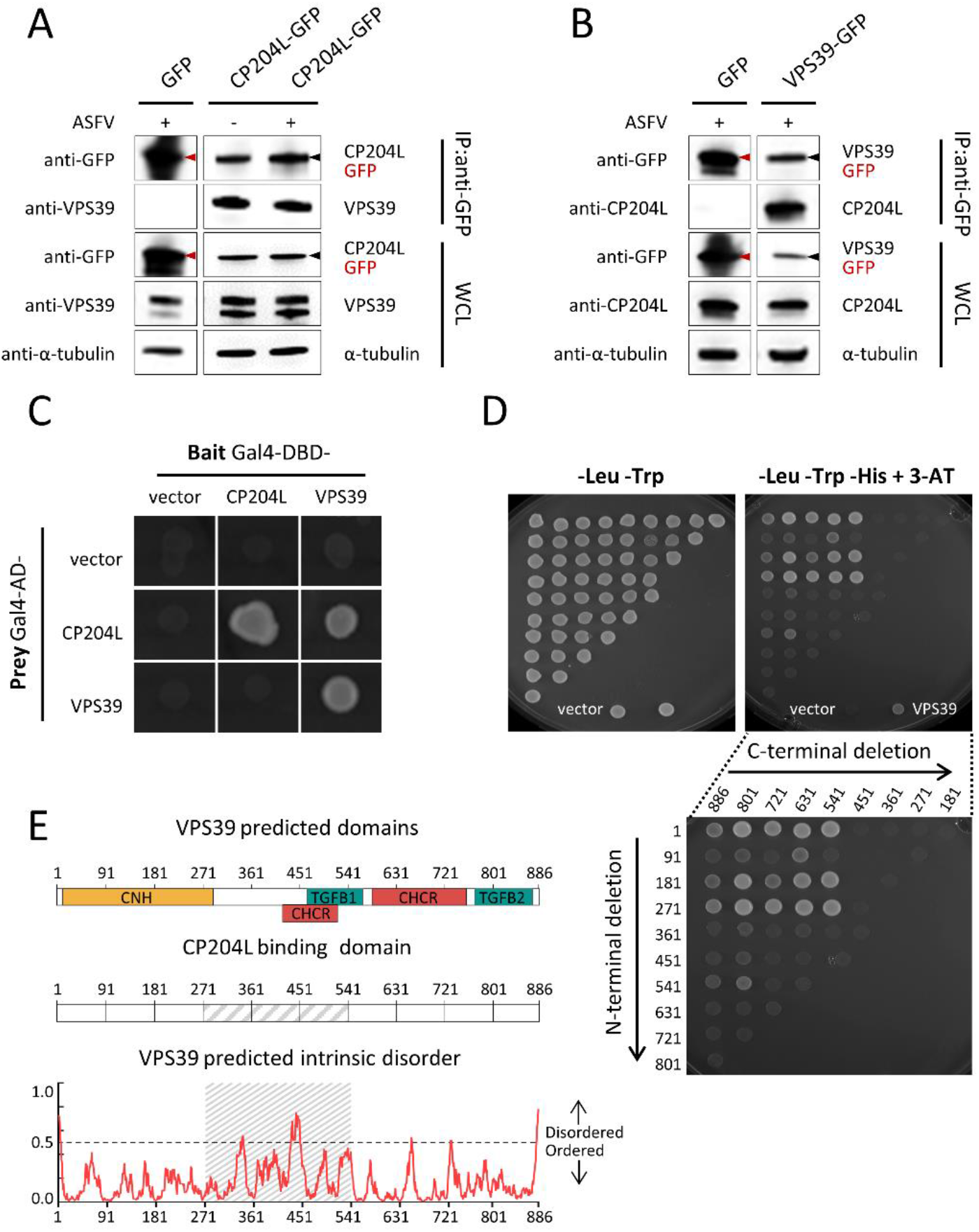
Interaction between ASFV CP204L and porcine VPS39. (A) Co-immunoprecipitation of GFP-tagged CP204L with endogenous VPS39. GFP was immunoprecipitated in lysates from mock and virus-infected cells stably expressing CP204L-GFP or GFP alone. Representative immunoblots of whole cell lysates (WCLs) and GFP-immunoprecipitates (IPs) are shown. α-tubulin was used as a loading control in WCLs. The black and red arrowheads indicate the bait proteins CP204L-GFP and GFP, respectively. (B) Reverse co-immunoprecipitation of VPS39-GFP with CP204L in virus-infected cells. A black arrowhead indicates bait protein VPS39-GFP. A red arrowhead indicates GFP used as a control. (C) Interaction between CP204L and VPS39 was tested in a yeast two-hybrid assay. CP204L and VPS39 were expressed as fusion proteins with the DBD or AD domain of Gal4. The empty vector is shown as a negative control. (D) Yeast two-hybrid mapping of CP204L-binding region in VPS39. Various combinations of VPS39 truncations and CP204L were co-transformed into the pPC86 vector. The transformants were spotted on control plates (-Leu -Trp) and selective plates (-Leu -Trp -His + 5mM 3-AT). Cotransfection with interacting protein fragments was indicatd by growth on the selective medium. Vertical and horizontal axes indicate the first and the last amino acid residues of each tested fragment, respectively. (E) The upper panel shows the predicted domain organization of porcineVPS39. CNH, citron homology domain; CHCR, clathrin heavy chain repeat; TGFB1 and TGFB2, transforming growth factor beta receptor-associated domain 1 and 2. The middle panel presents the CP204L-binding region between amino acids 271 and 541 of VPS39. The bottom panel shows the degree of order predicted by IUPred2A. The cut-off was set to a 0.5 probability score.

**Fig S3.**
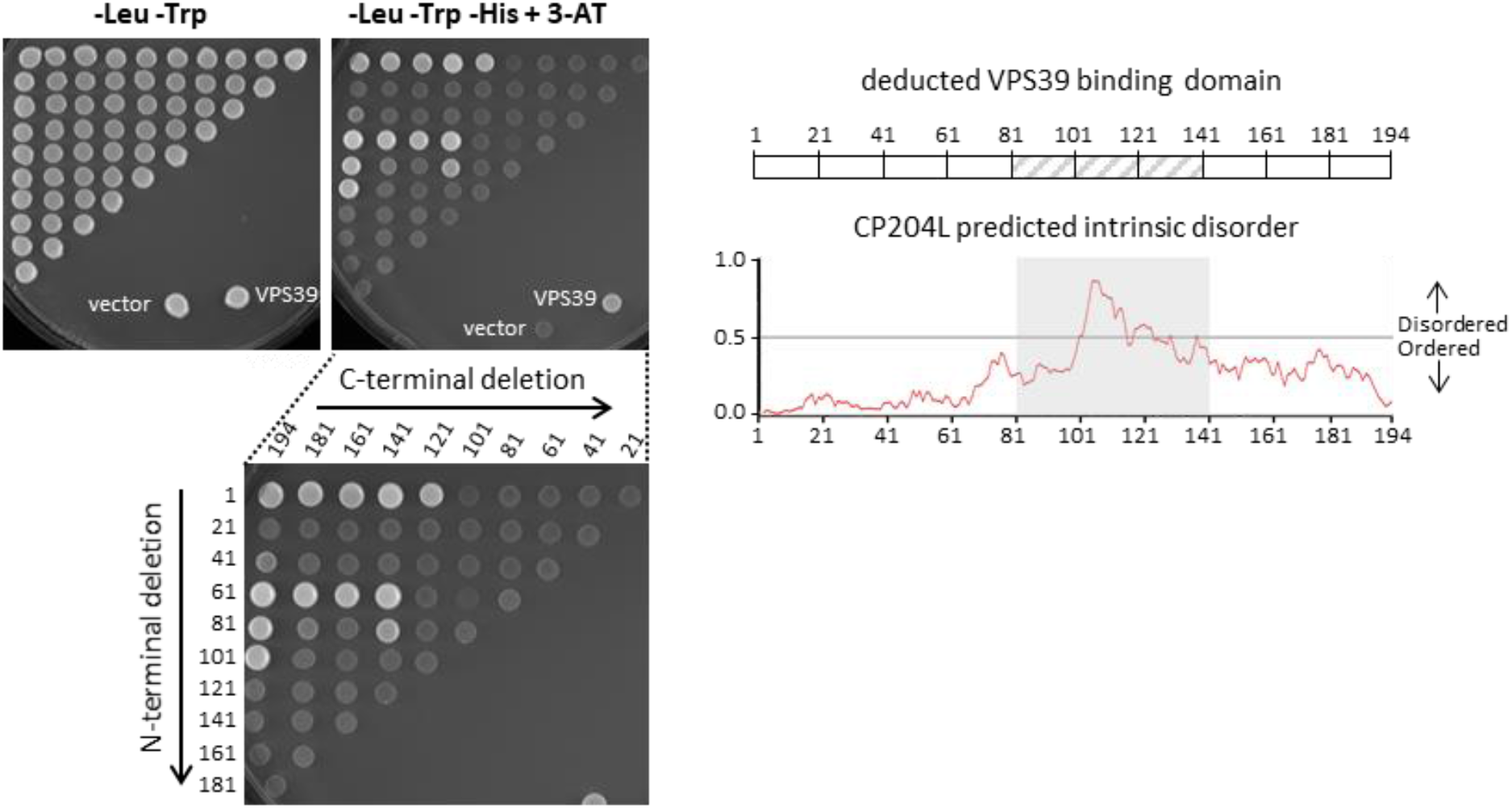
The deducted VPS39-binding region of CP204L The results of yeast two-hybrid mapping of VPS39-binding region in CP204L. Vertical and horizontal axes indicate the first and the last amino acid residues of each fragment tested, respectively. The right panels show the deducted VPS39-binding region of CP204L and the degree of order predicted by IUPred2A. The cut-off was set to a 0.5 probability score. The binding region of VPS39 was deduced to be localized between amino acids 81 and 141 of CP204L.

### CP204L and VPS39 colocalize and aggregate at the virus factory

We next performed colocalization analyses by confocal immunofluorescence microscopy. To improve the detection of the host protein, we used WSL cells stably expressing GFP-tagged VPS39. Cells were mock- or virus-infected and stained with anti-CP204L antibody after 24 hours. In the context of ASFV infection, both proteins colocalized in large cytoplasmic aggregates (Fig. 3A), whereas no VPS39 aggregates were detected in the absence of ASFV. A transfection experiment with a CP204L expression plasmid was performed to exclude the possibility of other virus proteins or infection-dependent mechanisms mediating the formation of VPS39-CP204L aggregates. This experiment confirmed that transiently expressed CP204L alone promotes VPS39 aggregation (Fig. 3B). Interestingly, the VPS39-CP204L complex formed large aggregates in the perinuclear region with smaller and more uniformly distributed granules. This observation led to the question of whether CP204L is targeted to a specific cellular structure in virus-infected cells in the absence of overexpressed VPS39. To address this question, WSL cells were infected with ASFV, and CP204L was visualized by indirect immunofluorescence 24 hours postinfection. We noticed that CP204L exhibited distinct distribution patterns in infected cells. In addition to an even distribution in the cytoplasm, which has been described in previous studies (24, 17), it accumulated at the site of virus factories that are enclosed in characteristic vimentin cages (Fig. 3C). CP204L accumulations were significantly larger in early virus factories than in the late ones (Fig. S4). Z-stack sectioning and imaging of infected cells showed the accumulation of CP204L around the sites of virus DNA replication (Fig. 3D). As expected, we could also confirm that VPS39 aggregates together with CP204L at the periphery of the ASFV replication sites (Fig. 3E).

**Fig 3.**
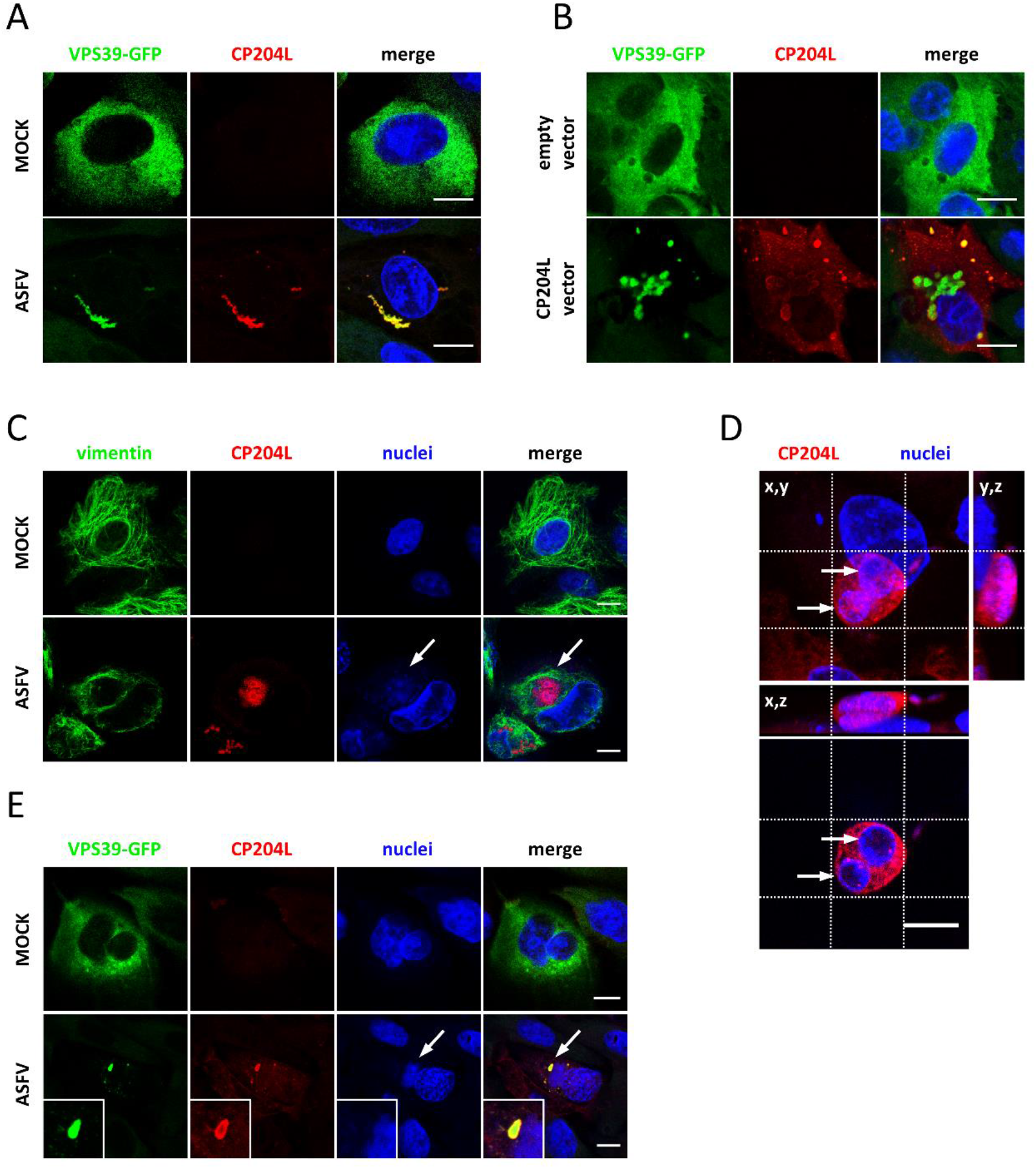
During infection CP204L and VPS39 form aggregates and localize to the periphery of ASFV replication sites. Colocalization and aggregate formation between CP204L and VPS39 in WSL-VPS39-GFP cells (A) 24 hours after ASFV-infection and (B) CP204L plasmid transfection. Empty vector and mock-infected cells were used as negative controls. (C) Subcellular localization of CP204L in infected WSL cells. Indirect immunofluorescence shows localization of CP204L (red) to virus replication sites (white arrows) surrounded by a vimentin cage (green). (D) Representative image of a WSL cell infected with ASFV (24 hours postinfection). The image shows a z-stack projection (64 slices across 12.8 μm) of the cell nucleus and virus factories (blue) and the CP204L protein (red). The crosshairs were positioned to indicate virus DNA. The bottom panel shows the cross-section through the replication compartment and the CP204L accumulation at its periphery. (E) Protein VPS39 (green) and CP204L (red) colocalization at the periphery of the virus replication site in infected WSL-VPS39-GFP cells. Hoechst dye was used to stain cellular and virus DNA (blue). White arrows indicate virus factories. Scale bars, 10 μm.

**Fig S4.**
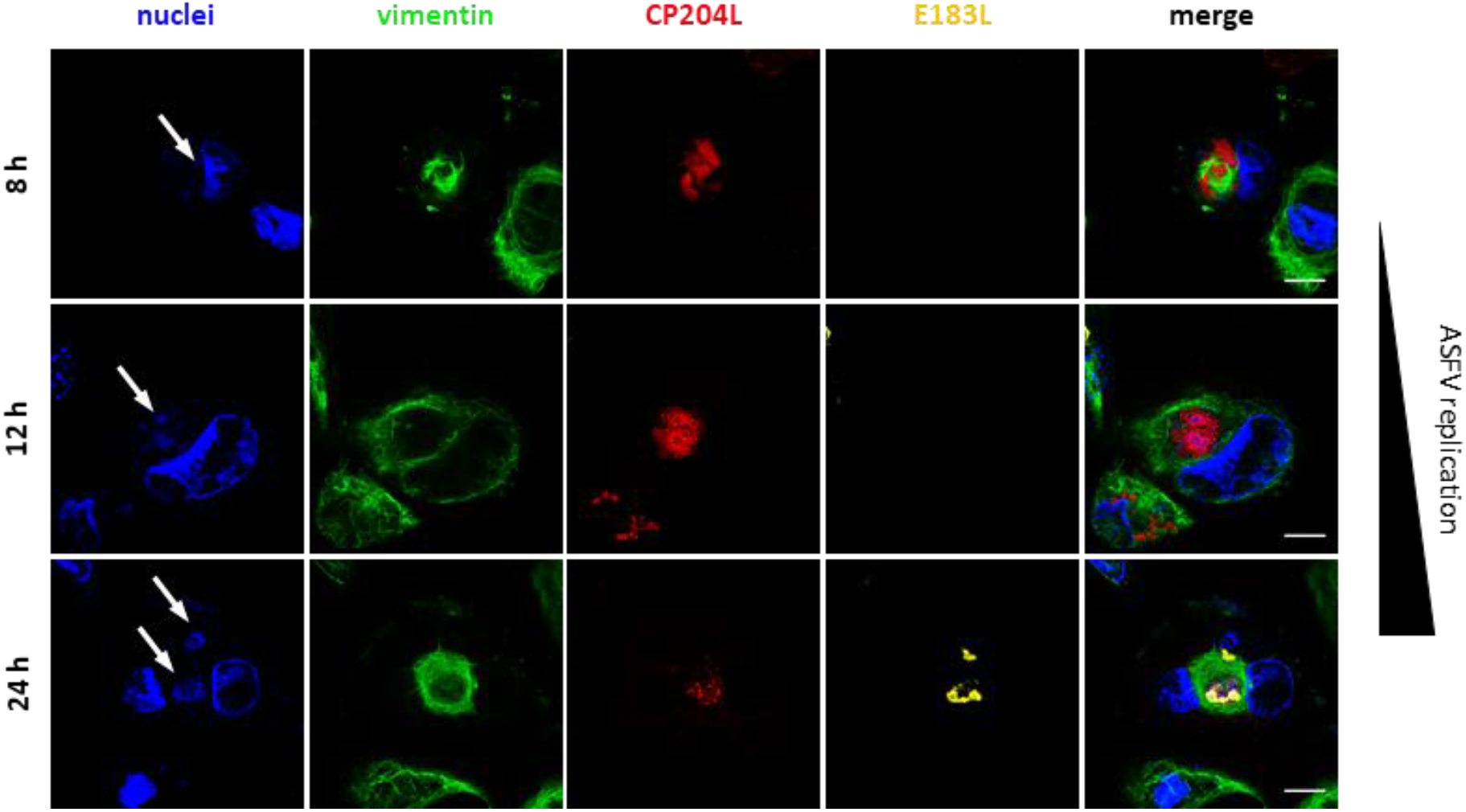
CP204L accumulates at the site of ASFV replication early during infection. Virus factories in WSL cells infected with ASFV were monitored by immunofluorescence at 8, 12, and 24 hours postinfection. Representative images show the accumulation of CP204L (red) in early VF and dispersion from late VF, which are marked by the presence of late virus protein E183L (P54) (yellow). Virus factories (white arrows) are labeled by DNA staining (blue) and vimentin cages (green). Scale bars, 10 μm.

### CP204L blocks VPS39 interaction with the HOPS complex and promotes lysosomal clustering

Having established that both proteins, CP204L and VPS39, interact and aggregate at the site of virus replication, we sought to determine whether this interaction could impair VPS39 function in late endosomal trafficking (33). The WSL cells expressing VPS39-GFP were transfected with a vector for CP204L expression or infected with ASFV. Cells expressing an empty vector were used as a control. After 24 hours, the colocalization of VPS39 with the lysosome-associated membrane protein 1 (LAMP1) was examined by immunofluorescence microscopy. Control cells exhibited the characteristic cytoplasmic and juxtanuclear distribution of lysosomes, and LAMP1 puncta colocalized with VPS39 (Fig. 4A). In the cells expressing CP204L after transfection or after infection, VPS39 aggregates were largely separated from LAMP1-marked lysosomes, which concentrated in the area near the nucleus. When CP204L was present, colocalization between the LAMP1-marked lysosomes and VPS39 was significantly reduced (Fig. 4B). Similarly, CP204L expression altered the colocalization between VPS39 and Rab7-marked late endosomes (Fig. S5). Moreover, the expression of CP204L was accompanied by reduced numbers of LAMP1-positive puncta and a significant increase in the size of accumulated lysosomes, suggesting lysosomal enlargement through coalescence (Fig. 4C). Finally, to test if the interaction with CP204L prevents the recruitment of VPS39 to the HOPS complex, we evaluated the colocalization of VPS39 with the subunit VPS11, which anchors VPS39 to the HOPS complex (34). Unlike in uninfected cells, no colocalization between VPS39 and VPS11 was observed in virus-infected cells (Fig. 4D). However, a clear colocalization between viral DNA (Hoechst staining) and VPS11 was detected, suggesting that VPS11 is targeted to ASFV replication site independent of VPS39. Together, these observations indicate that CP204L inhibits the integration of VPS39 into the HOPS complex, thus interfering with VPS39 function to associate with endosomal and lysosomal membranes and leading to the clustering of lysosomes in the infected cells.

**Fig 4.**
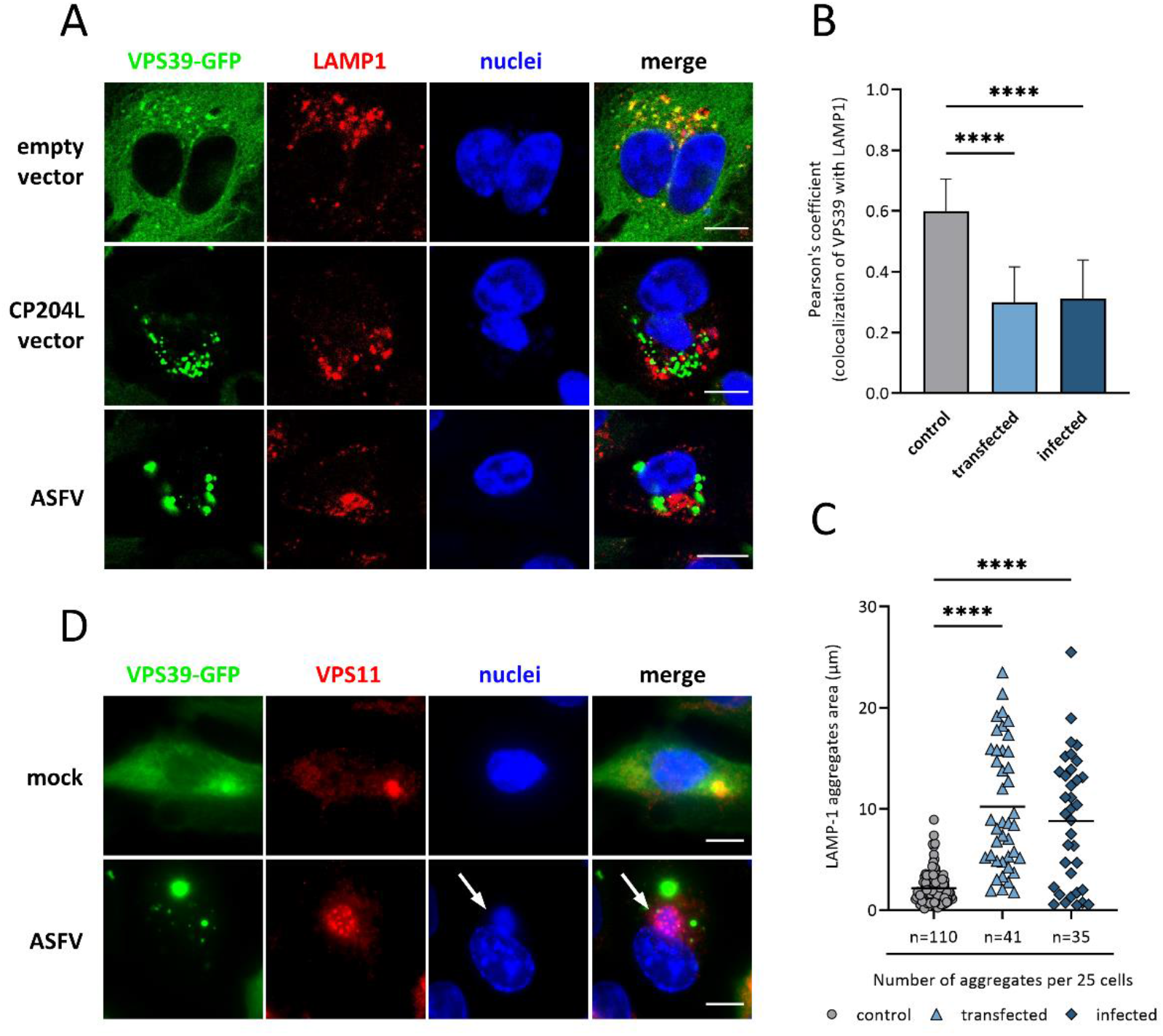
CP204L promotes lysosomal clustering and blocks VPS39 integration into the HOPS complex. (A) WSL cells stably expressing VPS39-GFP were analyzed by confocal fluorescence microscopy after transfection with a CP204L expression plasmid or infection with ASFV. Cells were stained for the lysosomal marker anti-LAMP1. (B) Quantification of the colocalization of VPS39 with lysosomes labeled by LAMP1. Pearson’s coefficient (mean ± SEM) from 25 cells in each group. (C) Quantification of the number and area of lysosomal and late endosomal aggregates per cell (n = 25 cells in each group). **** *p* < 0.0001. (D) VPS11 staining shows a loss of colocalization between VPS39 and VPS11 in virus-infected cells. White arrows indicate virus factories. Scale bars, 10μm.

**Fig S5.**
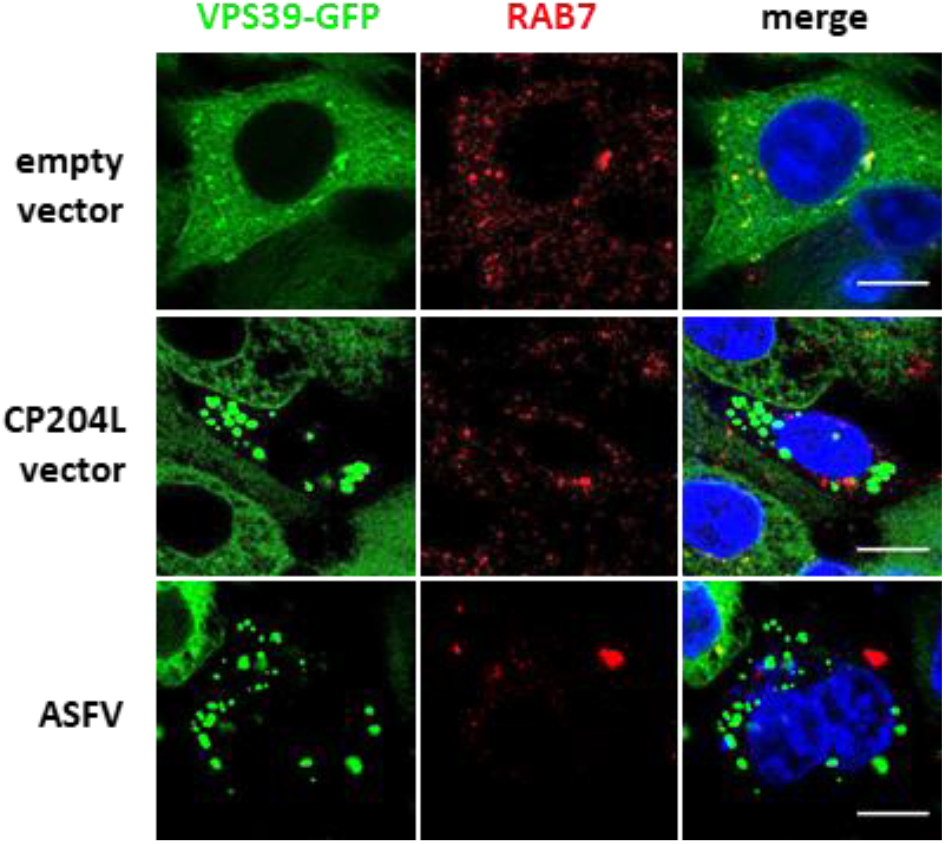
CP204L blocks VPS39 targeting to late endosomes. WSL cells stably expressing VPS39-GFP were observed by fluorescence confocal microscopy after transfection with CP204L expression construct or infection with ASFV. Cells were stained with anti-Rab7 antibody, a late endosomal marker. Scale bars, 10 μm.

### ASFV replication and protein synthesis are delayed in cells lacking VPS39

Next, we studied the role of the CP204L-VPS39 interaction in establishing the ASFV infection. CRISPR-Cas9 technology was used to generate a VPS39 knockout (KO) in the WSL cell line. The *VPS39* gene of the selected WSL cell clones exhibited a single nucleotide (A) insertion inducing a frameshift behind codon 202 (VPS39-KO1) and 101 (VPS39-KO2), leading to premature termination at positions 224 and 103, respectively. The absence of VPS39 was confirmed in the VPS39-KO cells by mass spectrometry, where no peptides corresponding to the VPS39 protein were detected in VPS39-KO1 and VPS39-KO2 (Fig. 5A). Importantly, VPS39 deficiency in WSL cells did not reduce the cell viability when compared to empty vector control cells (CTRL-KO) (Fig. 5B). Next, KO cells were infected with ASFV at the multiplicity of infection (MOI) of 1 and incubated for 8, 24 or 48 hours. Progeny virus titers were reduced by 1.5-log10 in VPS39-KO compared to control cell supernatant 24 hours after infection (Fig. 5C). However, final ASFV titers in VPS39-KO cells at 72 hours were not different from those in WT or CTRL-KO cells, suggesting that ASFV can replicate in the absence of VPS39, but with delayed kinetics. Additionally, the lysates of KO cells infected with ASFV for 8, 24, or 48 hours were harvested for Western blotting to examine the expression of early (CP204L) and late (B646L) virus proteins. While the expression of capsid protein B646L (P72) was similar in CTRL-KO and VPS39-KO cells, CP204L expression appeared to be delayed in cells lacking VPS39 (Fig. 5D). This observation raised the question of whether the synthesis of other early virus proteins expressed before the onset of virus DNA replication is also affected by the lack of VPS39. To answer this question, we first analyzed the changes in virus protein levels in CTRL-KO and VPS39-KO cells 8 hours after ASFV infection by quantitative MS. Among 46 virus proteins identified in all three cell lines, 22 were significantly down-regulated in cells lacking VPS39 (Fig. 6A). In particular, we observed a decrease in levels of virus proteins implicated in RNA transcription (i.e., ASFV RNA polymerase subunits: D205R, EP1242L, I243L, and D339L) and DNA replication (i.e., ribonucleotide reductase: F334L, F778R, and dUTPase E165R) (Fig. 6B). We further analyzed the dynamics of viral protein synthesis in VPS39-KO cells across time-points indicated (Fig. 6C; Table S5). As expected based on ASFV growth kinetic, most profound changes were observed at 8 hours after infection, whereas at later stages of infection, the levels of viral proteins in VPS39-KO cells stabilized and were comparable with those observed in CTRL-KO cells. These results suggest that although a loss of VPS39 markedly delays the synthesis of CP204L and other early proteins, VPS39 is not essential, and ASFV replication can proceed in its absence.

**Fig 5.**
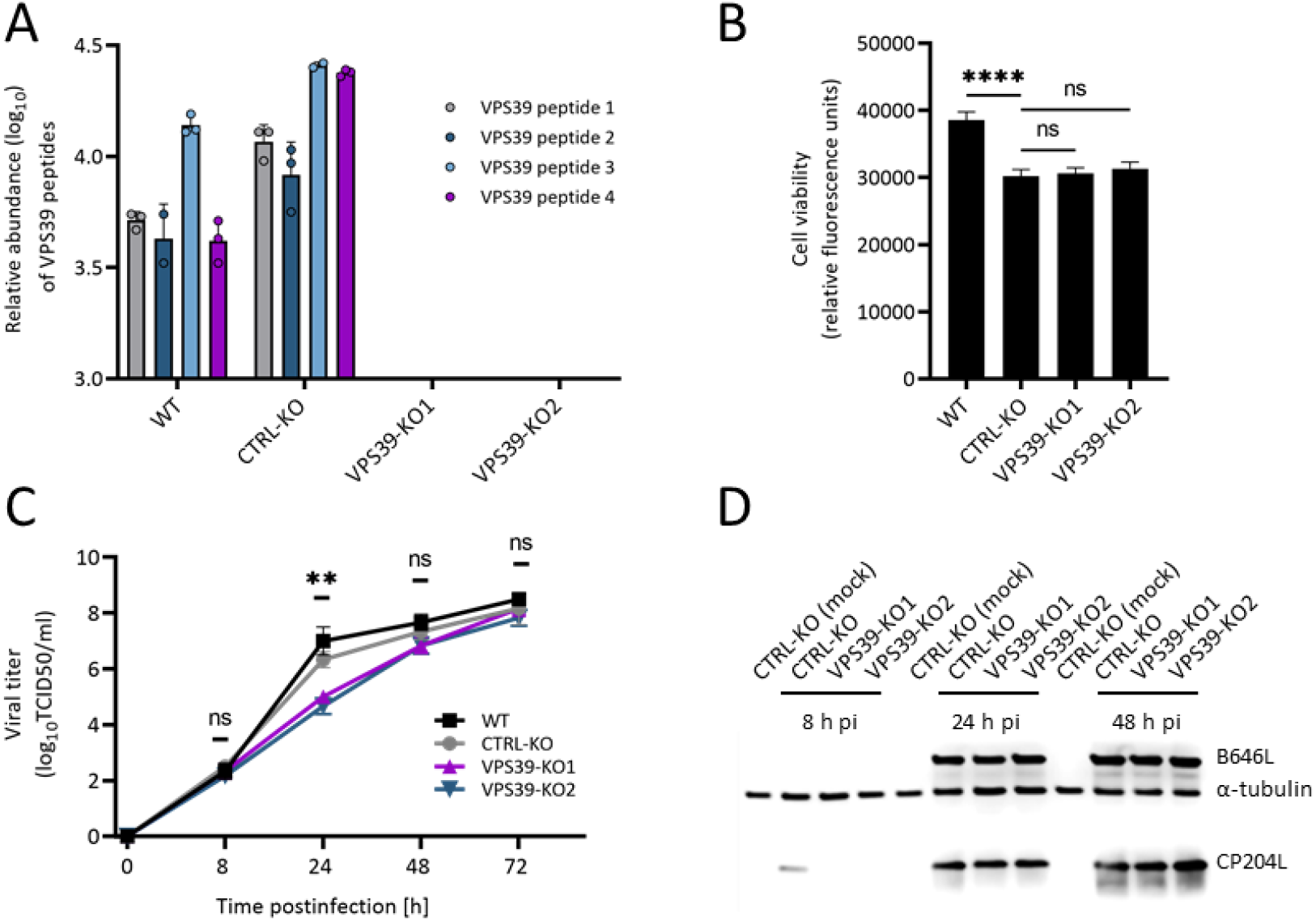
Absence of VPS39 delays ASFV growth and CP204L synthesis. (A) The absence of VPS39 in WSL KO cells was confirmed by mass spectrometry (see supplemental material Table S5). VPS39 was identified based on four unique peptides in wild-type (WT) WSL cells and control knockout (CTRL-KO) cells. In VPS39-KO cells, no peptides of VPS39 were detected. The log10 relative abundance is presented for each peptide identified from three independent experiments. (B) Cell viability of WSL WT, CTRL-KO, and VPS39-KO cell lines was quantitated by resazurin-based assay (PrestoBlue™). **** *p* < 0.0001. (C) Growth curves of ASFV after infection of VPS39-KO and control cells at an MOI of 1 (n = 3 wells/cell line/time point). The culture medium was collected at the indicated times, and the yields of the cell-free virus were expressed as 50% tissue culture infectious doses (TCID50)/ml and were plotted as means of results from three independent replicates and standard deviations. ** *p* < 0.01. (D) Expression of CP204L and B646L virus proteins in VPS39-KO and CTRL-KO cells. Mock-infected CTRL-KO cells were used as control. The α-tubulin specific monoclonal antibody was used as a loading control.

**Fig 6.**
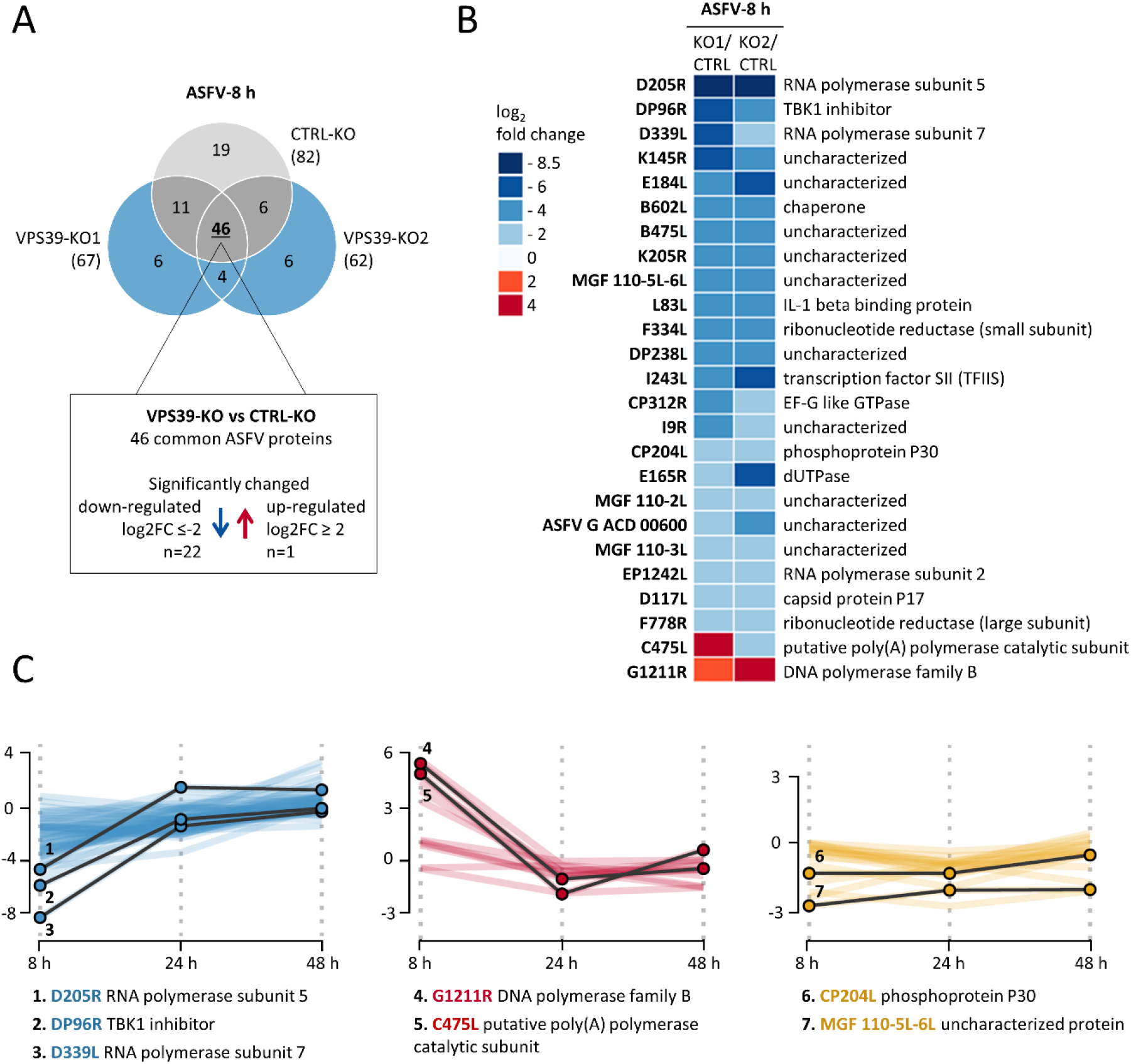
Loss of VPS39 affects the synthesis of ASFV proteins during the early phase of infection. (A) Overview of identified ASFV proteins 8 hours after infection in CTRL-KO and VPS39-KO cells. The number of proteins exhibiting reduced or increased amounts in VPS39-KO cells was determined. (B) A Heatmap of viral proteins identified at 8 hours depicts changes in protein expression levels between CTRL-KO and VPS39-KO cells. (C) Line plots showing fold changes in ASFV proteins expressed in VPS39-KO cells compared to control. Three clusters with similar profiles of protein were distinguished: down-regulated (or unchanged) at 8 h and increasing over time (blue); up-regulated (or unchanged) at 8 h and decreasing over time (red); continuously down-regulated during infection in VPS39-KO cells (yellow). Selected ASFV protein profiles are highlighted as solid black lines.

**Table S5**. Proteomic validation of VPS39 knockout and analysis of changes of ASFV protein levels between VPS39-KO and CTRL-KO cells at 8, 24, and 48 hours postinfection.

## Discussion

The protein interaction networks yield critical insights into the virus-host interrelationships and mechanisms of viral protein function. In this study, the protein interaction network derived from AP-MS data provided new information about the molecular hijacking strategies of ASFV and the role of virus protein CP204L. We found it particularly interesting that CP204L establishes a rich network of interactions with host vesicle trafficking proteins.

Vesicular trafficking is a directed cellular process of transporting cargo to target locations inside or outside the cell. It is, therefore, not surprising that viruses usurp this pathway to achieve efficient transport within the host cell. However, the role of host trafficking vesicles extends beyond virus movement, as the formation of virus-modified endosomal membranes supports viral replication, reported especially for RNA viruses (reviewed in (35)). It has been proposed that the cholesterol-rich endosomes can provide lipids for ASFV nucleic acid replication and virus assembly, structural scaffolding for viral factories, and protection from antiviral host responses (19, 36). Yet the precise contributions of cellular factors and mechanisms underlying endosomal recruitment and accumulation at the ASFV replication site remain largely enigmatic. Our study sheds new light on the mechanism of endosomal membrane redistribution during ASFV infection by identifying novel viral and cellular proteins involved in this process.

The important initial observation of this study was that the CP204L-host interactome is significantly enriched in proteins associated with vesicle transport and mitochondria (Fig. 1D). Within these two functional groups, VPS39, a subunit of the vacuolar HOPS tethering complex, and TOMM22, a subunit of the translocase of the outer membrane (TOM) mitochondrial complex, were the most abundant host interactors of ASFV CP204L (Fig. 1E). Interestingly, the interaction between the TOM complex and VPS39 was previously reported to be essential for the formation of membrane contact sites (MCSs) between cellular vesicles (e.g., endosomes, lysosomes) and mitochondria (37, 38). Only recently, MCSs have emerged as important host cell structures that enable viruses to reorganize the cellular membranes and channel cell lipids to the growing replication centers (39, 40). Therefore, it is tempting to speculate that the interaction between VPS39, TOMM22, and CP204L could play a role in establishing MCSs in ASFV-infected cells. However, to focus specifically on the mechanism by which ASFV exploits the endocytic pathway, this study solely investigates the interaction between VPS39 and CP204L.

We showed that CP204L directly and specifically binds VPS39, and the binding site could be mapped to amino acids 271 and 541 of VPS39 (Fig. 2). We could also confirm the formation of homo-oligomer for both proteins, CP204L (41) and VPS39 (42). Oligomerized CP204L is present in infected cells mostly as hexamers, and the existing dimers are suggested to serve as assembly units for the final oligomerization (41). While oligomerization is a common feature of many viral proteins, the specific mechanism and role of the CP204L oligomerization are not clear. In this context it is interesting that e.g., Ebola virus VP40 (43), influenza virus matrix protein M1 (44), and Dengue virus NS1 (45) undergo oligomerization when binding to cellular membranes. The formation of oligomers and the presence of intrinsically disordered regions in viral proteins enhance diverse interactions with host proteins, resulting in protein multifunctionality (46). Previously, Hernaez et al. (26) showed that a small portion of CP204L is present in the nucleus, interacting with heterogeneous nuclear ribonucleoprotein K (hnRNP-K), causing its retention in the nucleus. To date, CP204L was described as predominantly cytoplasmic ASFV protein. In this study, we show that CP204L localizes to VFs, specific intracellular sites of virus replication (Fig. 3). Localization of CP204L varies during infection, and its accumulation to the VFs occurs at early times, before ASFV late protein synthesis. Interestingly, immunofluorescence analysis revealed that CP204L is localized within VFs and is clearly separated from viral DNA, suggesting its possible role in viral protein transcription and translation rather than virus replication. Also, the fact that a large number of ASFV proteins were found to coimmunoprecipitate with CP204L (Fig. 1E) supports its possible role in the process of viral protein synthesis. On the other hand, it is also possible that CP204L interacts with proteins of membrane-bound organelles such as mitochondria, endoplasmic reticulum, or endosomes/lysosomes, which are all recruited to the area next to the virus replication site.

Recent findings of Miao et al. (47) and Zhang et al. (48) report direct interaction of SARS-CoV-2 protein ORF3a with the HOPS component VPS39. In this case, late endosome-localized ORF3a sequesters VPS39 and, consequently, inhibits autophagy by blocking the fusion of autophagosomes with lysosomes. Our results indicate that the mechanism by which ASFV protein CP204L interacts with VPS39 differs from the one used by SARS-CoV-2. Firstly, CP204L prevents VPS39 from binding to endosomal/lysosomal membranes (Fig. 4A, S4A). Secondly, CP204L expression inhibits VPS39 binding to VPS11 and its integration into the HOPS complex (Fig. 4D), while SARS-CoV-2 ORF3a expression does not affect the formation of the VPS39-containing HOPS complex (47). Thirdly, the expression of CP204L leads to clustering of lysosomes (Fig. 4C) and accumulation of CP204L-VPS39 aggregates near ASFV VFs (Fig. 3E).

CP204L interacts with VPS39 within its clathrin heavy chain region (CHCR), which is required for association with endosomal membranes via binding to RAB7 interacting lysosomal protein (RILP), and homooligomerization of VPS39 (49). RAB7 was already shown to be essential for ASFV replication (13). Therefore, the significant decrease in VPS39-lysosomes colocalization during ASFV infection can be explained by a simple competition model wherein highly abundant CP204L competes with host RILP for binding to VPS39 in the cytosol. The loss of the ability of VPS39 to bind VPS11 and, consequently, to form the HOPS complex in the presence of CP204L could be explained either by direct competition of CP204L and VPS11 for the same binding site on VPS39 or by an allosteric effect of the CP204L-VPS39 interaction on the VPS39 binding site for VPS11.

CP204L is essential for the virus, and the reduction of protein levels results in a strong suppression of viral replication (21). Although virus replication (Fig. 5) and protein synthesis (Fig. 6) were reduced in VPS39 deficient cells, especially early during infection, this effect did not correlate with the four orders of magnitude reduction observed by CP204L inhibition. This observation suggests that CP204L engages in several distinct functions important for virus replication, one of which involves the interaction with VPS39.

Nevertheless, we propose that the hijacking of VPS39 by CP204L may have two possible functions during ASFV infection. First, by inhibiting VPS39 association with endosomes and lysosomes, CP204L may impair homotypic and heterotypic fusion of vesicles, thus, protecting endocytosed viruses from degradation. Second, by CP204L-induced dissociation of VPS39 from the HOPS complex and endosomal membranes, VPS39 gains the ability to engage in membrane contact sites formation (37), playing a role in the biogenesis of ASFV VFs. In this case, the loss of VPS39 could affect the ASFV membrane synthesis and virus assembly, thus leading to the observed delay in virus replication.

In summary, based on our results and previous studies, we conclude that CP204L is a multifunctional protein that directly interacts with the VPS39 subunit of the HOPS complex and is involved in endosomal trafficking. CP204L exists in multiple oligomeric forms, undergoes phosphorylation, and localizes to the cytoplasm, nucleus, and virus factory.

## Materials and methods

### Cells and virus

The wild boar lung-derived cells (named WSL throughout the text) (50), supplied by Friedrich-Loeffler-Institut Biobank (Catalog number CCLV-RIE 0379), were maintained at 37°C with 5% CO_2_ in Iscove′s modified Dulbecco’s medium (DMEM) mixed with Ham’s F-12 nutrient mix (1:1; *v*/*v*) supplemented with 10% fetal bovine serum (FBS). ASFV (Armenia/07 isolate) was adapted by serial passaging to more efficient replication in WSL cells. Passage 20 stocks were generated as described previously (21).

### ASFV *in vitro* infection

All experiments with ASFV were performed in a biocontainment facility fulfilling the safety requirements for ASF laboratories and animal facilities (Commission Decision 2003/422/EC, Chapter VIII). For infection experiments, WSL cell monolayers were inoculated with ASFV stock dilutions at MOI of 2 PFU/cell, and supernatants collected from uninfected cells were used for the mock-infected controls. After inoculation, cells were centrifuged for 1 h at 600 x *g* and 37°C. Next, cells were washed three times with phosphate-buffered saline (PBS), replenished with a medium containing 5% FBS, and incubated at 37°C with 5% CO_2_. Supernatants were harvested at appropriate times, and progeny virus titers were determined as TCID50/ml (51) on WSL cells.

### DNA transfection

WSL cells were transiently transfected with a CP204L, a VPS39, or an empty vector using K2 Multiplier and K2 transfection reagent (Biontex) following the manufacturer’s instructions. Stable cell lines were generated by transient DNA transfection of WSL cells with plasmids coding GFP, CP204L-GFP, or VPS39-GFP. Three days after transfection, cells were trypsinized, seeded into 96 well plates, and maintained in a medium containing 500 μg/ml G418 (Corning). Single resistant and fluorescent cell clones detected after 2 to 3 weeks were further propagated and validated for expression of GFP fusion protein by immunoblotting.

### Plasmids constructs

The plasmid pUC-BaKJCAG-CP204Lsyn used for CP204L expression was previously described (52). pGFP-N1 plasmid (Clontech, GenBank accession # U55762) was used for GFP expression. Plasmid for expression of CP204L-GFP was created by amplification of CP204L from pUC-BaKJCAG-CP204Lsyn plasmid and insertion into pGFP-N1. The porcine VPS39 (isoform X3, GenBank accession # XP_013848582) was generated by gene synthesis (GeneArt, ThermoFisher Scientific) and recloned to pGFP-N1. Control plasmid (empty vector) was obtained by GFP deletion from pGFP-N1, resulting in pΔGFP-N1. All plasmid constructs were verified by DNA sequencing. The supplemental material includes a description of the cloning procedure and a list of primers and gene sequences (Text S1).

### Affinity purification and mass spectrometry

GFP-tagged target proteins and interacting proteins were affinity purified using GFP-trap agarose beads (Chromotek). Proteins were processed, trypsin digested, and concentrated for LC-MS/MS as described in the supplemental material (Text S1). Digested peptide mixtures were analyzed by LC-MS/MS on a timsTOF Pro (Bruker Daltonik), which was coupled online to a nanoElute nanoflow liquid chromatography system (Bruker Daltonik) via a CaptiveSpray nano-electrospray ion source. Data was analysed with MaxQuant (v.2.0.2.0) (53) and Perseus software (v.2.0.3.0) (54). Detailed information on sample preparation and analysis is provided in the supplemental material (Text S1).

### GO overrepresentation and network analysis

The obtained datasets for the CP204L interactome in mock and ASFV-infected cells were tested for enrichment of Gene Ontology (GO) biological process terms (55). Porcine genes were assigned to their corresponding human orthologues using the R package gprofiler2 (v.0.2.1) (56). The overrepresentation analysis was performed using the enricher function of clusterProfiler (v.4.2.2) (57) package in R with default parameters. Significant GO terms (adjusted *p*-value < 0.01) were identified and further clustered based on their semantic similarity using the R package rrvgo (v.1.6.0) (58). Selected preys were manually curated, and the network diagram was plotted using Cytoscape (v.3.7.2) (59).

### Immunoblotting

Beads and whole cell lysates were boiled in immunoprecipitation (IP) buffer (50 mM Tris-HCl, pH 7.4, 150 mM NaCl, 1 mM MgCl_2_) supplemented with benzonase (25 U/ml, Sigma-Aldrich #E8263), 0.5% Nonidet P40 substitute (NP-40; Sigma-Aldrich #I8896) and cOmplete mini EDTA-free protease inhibitor cocktail (Roche, #04693159001)], resolved on SDS-PAGE gels (4-20% Mini-PROTEAN TGX Gels (Bio-Rad) (60), and transferred to the nitrocellulose membrane by semidry transfer (Trans-Blot Turbo; Bio-Rad Laboratories) (61). All membranes were blocked in 5% milk powder in Tris-buffered saline with 0.25% Tween 20 (TBST) and probed overnight with the indicated primary antibodies using appropriate dilutions. This was followed by three 10 min washes in TBST and by incubation with peroxidase-conjugated secondary antibodies diluted in TBST. After 1 h, membranes were washed as above, and protein bands were detected using the Clarity Western ECL substrate (Bio-Rad) and imaged on C-DiGit Blot Scanner (LI-COR) and analyzed by Image Studio Software (v.5.2).

### Immunofluorescence

Coverslips were fixed with 3.7 % formaldehyde in PBS for 60 min at room temperature and then washed 3 times for 10 min with PBS, permeabilized with 0.01% Triton X-100 in PBS for 15 min, and then blocked with PBS containing 10% FBS for 1 h. Coverslips were incubated with the primary antibody for one hour at 37°C and then with the secondary antibody for 1 hour at 37°C. Nuclei were stained for 15 min with 1 μg/ml Hoechst 33258 (Sigma-Aldrich) in PBS. After each step, the cells were repeatedly washed with PBS. Coverslips were then mounted on glass slides using. Images were acquired on a Leica DMI6000 TCS SP5 confocal laser scan microscope (63× objective) and were processed with the ImageJ software (v.1.52a) (62).

### Antibodies

The ASFV CP204L, B646L, E183L (52), and A137R (unpublished) protein-specific rabbit antisera were used at dilutions of 1:20 000 for immunoblotting. The primary antibodies used for immunoblotting included rabbit anti-GFP (Chromotek), rabbit anti-VPS39 (PA5-21104; Thermo Fisher), mouse anti-tubulin (B-5-1-2; Sigma-Aldrich), and mouse anti-GAPDH (MCA4739; BioRad). The secondary antibodies used were peroxidase-conjugated goat anti-mouse and anti-rabbit IgG (Jackson ImmunoResearch). The additional primary antibodies used for immunofluorescence were mouse anti-vimentin (MA1-06908; Thermo Fisher), rabbit anti-RAB7 (PA5-52369; Thermo Fisher), mouse anti-LAMP-1 (MCA2315GA; BioRad), rabbit anti-VPS11 (PA5-21854; Thermo Scientific). The secondary antibodies were Alexa Fluor 647-conjugated goat anti-rabbit IgG (H+L) or goat anti-mouse IgG (H+L), respectively (Invitrogen).

### Yeast two-hybrid analysis of the protein interaction

To identify the CP204L binding site, we used the yeast two-hybrid system. First, forward and reverse PCR primers were designed along the VPS39 sequence every 270 and 240 nucleotides. These primer sequences were fused to specific tails allowing recombination in the Gal4-BD pDEST32 vector. The sequences of the specific tails were 5′-GAAGAGAGTAGTAACAAAGGTCAAAGACAGTTGACTGTATCGTCGAGG-3′ and 5′-CCGCGGTGGCGGCCGTTACTTACTTAGAGCTCGACGTCTTACTTA-3′. Matrix combinations of forward and reverse primers were used to amplify VPS39 fragments by PCR. As previously described (63), 10 ng of linearized pDEST32 empty vector was co-transformed with 3 μl of PCR product to achieve recombinational cloning by gap-repair in Y2H Gold yeast strain (Clontech) expressing AD-fused CP204L (pPC86 vector). Interactions between VPS39 fragments and CP204L were tested by plating yeast cells on a selective medium lacking leucine, tryptophan, and histidine and supplemented with 5 mM of 3-amino-1,2,4-triazole (3-AT).

### Generation of CRISPR-Cas9 knockout cell lines

To generate VPS39 knockout cells, suitable CRISPR/Cas9 target sites were identified within the first 5’-terminal exons present in all predicted mRNA splice variants of the VPS39 gene (GenBank accession # NC_010443), and corresponding oligonucleotides were synthesized (Eurofins Genomics). The complementary oligonucleotide pairs VPS39porc-gR3F and VPS39porc-gR3F, as well as VPS39porc-gR4F and VPS39porc-gR4F (see Text S1), were phosphorylated and cloned into the BpiI-digested and dephosphorylated single guide RNA (sgRNA) and Cas9 nuclease expression vector pX330A-1×4neoR (unpublished results). After verifying the correct sequence insertions, the resulting plasmids pX330-VPS39gR3neoR and pX330-VPS39gR4neoR were used to transfect WSL cells, and single geneticin-resistant cell clones were selected with a medium containing 500 μg/ml G418. The clones were propagated, and the VPS39 knockout was verified as described in the supplemental material (Text S1). Similarly, a single cell clone was generated with an empty Cas9 nuclease expression vector pX330A-1×4neoR as a control (CTRL-KO).

## Data availability

All mass spectrometry raw data and MaxQuant output tables are deposited to the ProteomeXchange Consortium (http://proteomecentral.proteomexchange.org) via the PRIDE partner repository (64) and will be publicly available upon final publication (identifier PXD035695; accession for reviewers: Username; Password: 3Qh0Sega).

## Acknowledgments

We thank Biobank of the Friedrich-Loeffler-Institut for providing the WSL cell line. We also thank Barbara Bettin and Christin Milde for their technical assistance.

This work was supported by research grants from European Commission, Horizon 2020 Framework Programme European Union ERA-NET project “ASFVInt”, grant agreement 862605, and FLI’s ASFV Research Network. J.D. is supported by a Ph.D. fellowship from ANSES and INRAE.

T.C.M. and A.K. conceived and designed the study. S.B., T.C.M, and A.K acquired funding and resources. A.K. supervised the study. K.M.D., W.F., G.C, J.D., K.P. contributed to the methodology and investigation. K.D.M. wrote the first draft. K.D.M. and A.K. reviewed and edited the article. All authors read and approved the manuscript.

## References

1. Eustace Montgomery R. 1921. On A Form of Swine Fever Occurring in British East Africa (Kenya Colony). Journal of Comparative Pathology and Therapeutics 34:159–191. doi:10.1016/S0368-1742(21)80031-4.

2. Gabriel C, Blome S, Malogolovkin A, Parilov S, Kolbasov D, Teifke JP, Beer M. 2011. Characterization of African swine fever virus Caucasus isolate in European wild boars. Emerg Infect Dis 17:2342–2345. doi:10.3201/eid1712.110430.

3. Gallardo C, Soler A, Nieto R, Cano C, Pelayo V, Sánchez MA, Pridotkas G, Fernandez-Pinero J, Briones V, Arias M. 2017. Experimental Infection of Domestic Pigs with African Swine Fever Virus Lithuania 2014 Genotype II Field Isolate. Transbound Emerg Dis 64:300–304. doi:10.1111/tbed.12346.

4. You S, Liu T, Zhang M, Zhao X, Dong Y, Wu B, Wang Y, Li J, Wei X, Shi B. 2021. African swine fever outbreaks in China led to gross domestic product and economic losses. Nat Food 2:802–808. doi:10.1038/s43016-021-00362-1.

5. Wöhnke E, Fuchs W, Hartmann L, Blohm U, Blome S, Mettenleiter TC, Karger A. 2021. Comparison of the Proteomes of Porcine Macrophages and a Stable Porcine Cell Line after Infection with African Swine Fever Virus. Viruses 13. doi:10.3390/v13112198.

6. Keßler C, Forth JH, Keil GM, Mettenleiter TC, Blome S, Karger A. 2018. The intracellular proteome of African swine fever virus. Sci Rep 8:14714. doi:10.1038/s41598-018-32985-z.

7. Alejo A, Matamoros T, Guerra M, Andrés G. 2018. A Proteomic Atlas of the African Swine Fever Virus Particle. J Virol 92. doi:10.1128/JVI.01293-18.

8. Gómez-Villamandos JC, Hervás J, Méndez A, Carrasco L, Villeda CJ, Wilkinson PJ, Sierra MA. 1995. Ultrastructural study of the renal tubular system in acute experimental African swine fever: virus replication in glomerular mesangial cells and in the collecting ducts. Arch Virol 140:581–589. doi:10.1007/BF01718433.

9. Kleiboeker SB, Scoles GA, Burrage TG, Sur J. 1999. African swine fever virus replication in the midgut epithelium is required for infection of Ornithodoros ticks. J Virol 73:8587–8598. doi:10.1128/JVI.73.10.8587-8598.1999.

10. Hernaez B, Alonso C. 2010. Dynamin- and clathrin-dependent endocytosis in African swine fever virus entry. J Virol 84:2100–2109. doi:10.1128/JVI.01557-09.

11. Sánchez EG, Quintas A, Pérez-Núñez D, Nogal M, Barroso S, Carrascosa ÁL, Revilla Y. 2012. African swine fever virus uses macropinocytosis to enter host cells. PLoS Pathog 8:e1002754. doi:10.1371/journal.ppat.1002754.

12. Basta S, Gerber H, Schaub A, Summerfield A, McCullough KC. 2010. Cellular processes essential for African swine fever virus to infect and replicate in primary macrophages. Vet Microbiol 140:9–17. doi:10.1016/j.vetmic.2009.07.015.

13. Cuesta-Geijo MA, Galindo I, Hernáez B, Quetglas JI, Dalmau-Mena I, Alonso C. 2012. Endosomal maturation, Rab7 GTPase and phosphoinositides in African swine fever virus entry. PLoS One 7:e48853. doi:10.1371/journal.pone.0048853.

14. Alcamí A, Carrascosa AL, Viñuela E. 1989. The entry of African swine fever virus into Vero cells. Virology 171:68–75. doi:10.1016/0042-6822(89)90511-4.

15. Hernáez B, Guerra M, Salas ML, Andrés G. 2016. African Swine Fever Virus Undergoes Outer Envelope Disruption, Capsid Disassembly and Inner Envelope Fusion before Core Release from Multivesicular Endosomes. PLoS Pathog 12:e1005595. doi:10.1371/journal.ppat.1005595.

16. Castelló A, Quintas A, Sánchez EG, Sabina P, Nogal M, Carrasco L, Revilla Y. 2009. Regulation of host translational machinery by African swine fever virus. PLoS Pathog 5:e1000562. doi:10.1371/journal.ppat.1000562.

17. Stefanovic S, Windsor M, Nagata K-I, Inagaki M, Wileman T. 2005. Vimentin rearrangement during African swine fever virus infection involves retrograde transport along microtubules and phosphorylation of vimentin by calcium calmodulin kinase II. J Virol 79:11766–11775. doi:10.1128/JVI.79.18.11766-11775.2005.

18. Aicher S-M, Monaghan P, Netherton CL, Hawes PC. 2021. Unpicking the Secrets of African Swine Fever Viral Replication Sites. Viruses 13. doi:10.3390/v13010077.

19. Cuesta-Geijo MÁ, Barrado-Gil L, Galindo I, Muñoz-Moreno R, Alonso C. 2017. Redistribution of Endosomal Membranes to the African Swine Fever Virus Replication Site. Viruses 9. doi:10.3390/v9060133.

20. Alcaraz C, Diego M de, Pastor MJ, Escribano JM. 1990. Comparison of a radioimmunoprecipitation assay to immunoblotting and ELISA for detection of antibody to African swine fever virus. J Vet Diagn Invest 2:191–196. doi:10.1177/104063879000200307.

21. Hübner A, Petersen B, Keil GM, Niemann H, Mettenleiter TC, Fuchs W. 2018. Efficient inhibition of African swine fever virus replication by CRISPR/Cas9 targeting of the viral p30 gene (CP204L). Sci Rep 8:1449. doi:10.1038/s41598-018-19626-1.

22. Zheng Y, Li S, Li S-H, Yu S, Wang Q, Zhang K, Qu L, Sun Y, Bi Y, Tang F, Qiu H-J, Gao GF. 2022. Transcriptome profiling in swine macrophages infected with African swine fever virus at single-cell resolution. Proc Natl Acad Sci U S A 119:e2201288119. doi:10.1073/pnas.2201288119.

23. Cackett G, Matelska D, Sýkora M, Portugal R, Malecki M, Bähler J, Dixon L, Werner F. 2020. The African Swine Fever Virus Transcriptome. J Virol 94. doi:10.1128/JVI.00119-20.

24. Prados FJ, Viñuela E, Alcamí A. 1993. Sequence and characterization of the major early phosphoprotein p32 of African swine fever virus. J Virol 67:2475–2485. doi:10.1128/JVI.67.5.2475-2485.1993.

25. Afonso CL, Alcaraz C, Brun A, Sussman MD, Onisk DV, Escribano JM, Rock DL. 1992. Characterization of P30, a highly antigenic membrane and secreted protein of African Swine Fever Virus. Virology 189:368–373. doi:10.1016/0042-6822(92)90718-5.

26. Hernaez B, Escribano JM, Alonso C. 2008. African swine fever virus protein p30 interaction with heterogeneous nuclear ribonucleoprotein K (hnRNP-K) during infection. FEBS Lett 582:3275–3280. doi:10.1016/j.febslet.2008.08.031.

27. Chapman DAG, Darby AC, Da Silva M, Upton C, Radford AD, Dixon LK. 2011. Genomic analysis of highly virulent Georgia 2007/1 isolate of African swine fever virus. Emerg Infect Dis 17:599–605. doi:10.3201/eid1704.101283.

28. Rink J, Ghigo E, Kalaidzidis Y, Zerial M. 2005. Rab conversion as a mechanism of progression from early to late endosomes. Cell 122:735–749. doi:10.1016/j.cell.2005.06.043.

29. Jiang P, Nishimura T, Sakamaki Y, Itakura E, Hatta T, Natsume T, Mizushima N. 2014. The HOPS complex mediates autophagosome-lysosome fusion through interaction with syntaxin 17. Mol Biol Cell 25:1327–1337. doi:10.1091/mbc.E13-08-0447.

30. Takáts S, Pircs K, Nagy P, Varga Á, Kárpáti M, Hegedűs K, Kramer H, Kovács AL, Sass M, Juhász G. 2014. Interaction of the HOPS complex with Syntaxin 17 mediates autophagosome clearance in Drosophila. Mol Biol Cell 25:1338–1354. doi:10.1091/mbc.E13-08-0449.

31. Blum M, Chang H-Y, Chuguransky S, Grego T, Kandasaamy S, Mitchell A, Nuka G, Paysan-Lafosse T, Qureshi M, Raj S, Richardson L, Salazar GA, Williams L, Bork P, Bridge A, Gough J, Haft DH, Letunic I, Marchler-Bauer A, Mi H, Natale DA, Necci M, Orengo CA, Pandurangan AP, Rivoire C, Sigrist CJA, Sillitoe I, Thanki N, Thomas PD, Tosatto SCE, Wu CH, Bateman A, Finn RD. 2021. The InterPro protein families and domains database: 20 years on. Nucleic Acids Res 49:D344–D354. doi:10.1093/nar/gkaa977.

32. Mészáros B, Erdos G, Dosztányi Z. 2018. IUPred2A: context-dependent prediction of protein disorder as a function of redox state and protein binding. Nucleic Acids Res 46:W329–W337. doi:10.1093/nar/gky384.

33. Pols MS, Brink C ten, Gosavi P, Oorschot V, Klumperman J. 2013. The HOPS proteins hVps41 and hVps39 are required for homotypic and heterotypic late endosome fusion. Traffic 14:219–232.

34. Bröcker C, Kuhlee A, Gatsogiannis C, Balderhaar HJk, Hönscher C, Engelbrecht-Vandré S, Ungermann C, Raunser S. 2012. Molecular architecture of the multisubunit homotypic fusion and vacuole protein sorting (HOPS) tethering complex. Proc Natl Acad Sci U S A 109:1991– 1996. doi:10.1073/pnas.1117797109.

35. Miller S, Krijnse-Locker J. 2008. Modification of intracellular membrane structures for virus replication. Nat Rev Microbiol 6:363–374. doi:10.1038/nrmicro1890.

36. Galindo I, Cuesta-Geijo MÁ, Del Puerto A, Soriano E, Alonso C. 2019. Lipid Exchange Factors at Membrane Contact Sites in African Swine Fever Virus Infection. Viruses 11. doi:10.3390/v11030199.

37. González Montoro A, Auffarth K, Hönscher C, Bohnert M, Becker T, Warscheid B, Reggiori F, van der Laan M, Fröhlich F, Ungermann C. 2018. Vps39 Interacts with Tom40 to Establish One of Two Functionally Distinct Vacuole-Mitochondria Contact Sites. Dev Cell 45:621-636.e7. doi:10.1016/j.devcel.2018.05.011.

38. Jackson J, Wischhof L, Scifo E, Pellizzer A, Wang Y, Piazzesi A, Gentile D, Siddig S, Stork M, Hopkins CE, Händler K, Weis J, Roos A, Schultze JL, Nicotera P, Ehninger D, Bano D. 2022. SGPL1 stimulates VPS39 recruitment to the mitochondria in MICU1 deficient cells. Mol Metab 61:101503. doi:10.1016/j.molmet.2022.101503.

39. Laufman O, Perrino J, Andino R. 2019. Viral Generated Inter-Organelle Contacts Redirect Lipid Flux for Genome Replication. Cell 178:275-289.e16. doi:10.1016/j.cell.2019.05.030.

40. Wong LH, Edgar JR, Martello A, Ferguson BJ, Eden ER. 2021. Exploiting Connections for Viral Replication. Front Cell Dev Biol 9:640456. doi:10.3389/fcell.2021.640456.

41. Anders et al. 1993 Characterization of Two ASFV 220kDa Proteins A Precursor of the Major Structural Protein p150 and Oligomer of Phosphoprotein p32.

42. Caplan S, Hartnell LM, Aguilar RC, Naslavsky N, Bonifacino JS. 2001. Human Vam6p promotes lysosome clustering and fusion in vivo. J Cell Biol 154:109–122. doi:10.1083/jcb.200102142.

43. Scianimanico S, Schoehn G, Timmins J, Ruigrok RH, Klenk HD, Weissenhorn W. 2000. Membrane association induces a conformational change in the Ebola virus matrix protein. EMBO J 19:6732–6741. doi:10.1093/emboj/19.24.6732.

44. Hilsch M, Goldenbogen B, Sieben C, Höfer CT, Rabe JP, Klipp E, Herrmann A, Chiantia S. 2014. Influenza A matrix protein M1 multimerizes upon binding to lipid membranes. Biophys J 107:912–923. doi:10.1016/j.bpj.2014.06.042.

45. Gutsche I, Coulibaly F, Voss JE, Salmon J, d’Alayer J, Ermonval M, Larquet E, Charneau P, Krey T, Mégret F, Guittet E, Rey FA, Flamand M. 2011. Secreted dengue virus nonstructural protein NS1 is an atypical barrel-shaped high-density lipoprotein. Proc Natl Acad Sci U S A 108:8003–8008. doi:10.1073/pnas.1017338108.

46. Jayaraman B, Smith AM, Fernandes JD, Frankel AD. 2016. Oligomeric viral proteins: small in size, large in presence. Crit Rev Biochem Mol Biol 51:379–394. doi:10.1080/10409238.2016.1215406.

47. Miao G, Zhao H, Li Y, Ji M, Chen Y, Shi Y, Bi Y, Wang P, Zhang H. 2021. ORF3a of the COVID-19 virus SARS-CoV-2 blocks HOPS complex-mediated assembly of the SNARE complex required for autolysosome formation. Dev Cell 56:427-442.e5. doi:10.1016/j.devcel.2020.12.010.

48. Zhang Y, Sun H, Pei R, Mao B, Zhao Z, Li H, Lin Y, Lu K. 2021. The SARS-CoV-2 protein ORF3a inhibits fusion of autophagosomes with lysosomes. Cell Discov 7. doi:10.1038/s41421-021-00268-z.

49. van der Kant R, Jonker CTH, Wijdeven RH, Bakker J, Janssen L, Klumperman J, Neefjes J. 2015. Characterization of the Mammalian CORVET and HOPS Complexes and Their Modular Restructuring for Endosome Specificity. J Biol Chem 290:30280–30290. doi:10.1074/jbc.M115.688440.

50. Keil GM, Giesow K, Portugal R. 2014. A novel bromodeoxyuridine-resistant wild boar lung cell line facilitates generation of African swine fever virus recombinants. Arch Virol 159:2421–2428. doi:10.1007/s00705-014-2095-2.

51. Reed LJ, Muench H. 1938. A SIMPLE METHOD OF ESTIMATING FIFTY PER CENT ENDPOINTS12. American Journal of Epidemiology 27:493–497. doi:10.1093/oxfordjournals.aje.a118408.

52. Hübner A, Keil GM, Kabuuka T, Mettenleiter TC, Fuchs W. 2018. Efficient transgene insertion in a pseudorabies virus vector by CRISPR/Cas9 and marker rescue-enforced recombination. J Virol Methods 262:38–47. doi:10.1016/j.jviromet.2018.09.009.

53. Cox J, Mann M. 2008. MaxQuant enables high peptide identification rates, individualized p.p.b.-range mass accuracies and proteome-wide protein quantification. Nat Biotechnol 26:1367–1372. doi:10.1038/nbt.1511.

54. Tyanova S, Temu T, Sinitcyn P, Carlson A, Hein MY, Geiger T, Mann M, Cox J. 2016. The Perseus computational platform for comprehensive analysis of (prote)omics data. Nat Methods 13:731–740. doi:10.1038/nmeth.3901.

55. Ashburner M, Ball CA, Blake JA, Botstein D, Butler H, Cherry JM, Davis AP, Dolinski K, Dwight SS, Eppig JT, Harris MA, Hill DP, Issel-Tarver L, Kasarskis A, Lewis S, Matese JC, Richardson JE, Ringwald M, Rubin GM, Sherlock G. 2000. Gene ontology: tool for the unification of biology. The Gene Ontology Consortium. Nat Genet 25:25–29. doi:10.1038/75556.

56. Kolberg L, Raudvere U, Kuzmin I, Vilo J, Peterson H. 2020. gprofiler2 -- an R package for gene list functional enrichment analysis and namespace conversion toolset g:Profiler. F1000Res 9. doi:10.12688/f1000research.24956.2.

57. Wu T, Hu E, Xu S, Chen M, Guo P, Dai Z, Feng T, Zhou L, Tang W, Zhan L, Fu X, Liu S, Bo X, Yu G. 2021. clusterProfiler 4.0: A universal enrichment tool for interpreting omics data. Innovation (N Y) 2:100141. doi:10.1016/j.xinn.2021.100141.

58. Sergi Sayols. 2020. rrvgo. Bioconductor.

59. Shannon P, Markiel A, Ozier O, Baliga NS, Wang JT, Ramage D, Amin N, Schwikowski B, Ideker T. 2003. Cytoscape: a software environment for integrated models of biomolecular interaction networks. Genome Res 13:2498–2504. doi:10.1101/gr.1239303.

60. Laemmli UK. 1970. Cleavage of structural proteins during the assembly of the head of bacteriophage T4. Nature 227:680–685. doi:10.1038/227680a0.

61. Towbin H, Staehelin T, Gordon J. 1979. Electrophoretic transfer of proteins from polyacrylamide gels to nitrocellulose sheets: procedure and some applications. Proc Natl Acad Sci U S A 76:4350–4354. doi:10.1073/pnas.76.9.4350.

62. Schneider CA, Rasband WS, Eliceiri KW. 2012. NIH Image to ImageJ: 25 years of image analysis. Nat Methods 9:671–675. doi:10.1038/nmeth.2089.

63. Walhout AJ, Vidal M. 2001. High-throughput yeast two-hybrid assays for large-scale protein interaction mapping. Methods 24:297–306. doi:10.1006/meth.2001.1190.

64. Vizcaíno JA, Deutsch EW, Wang R, Csordas A, Reisinger F, Ríos D, Dianes JA, Sun Z, Farrah T, Bandeira N, Binz P-A, Xenarios I, Eisenacher M, Mayer G, Gatto L, Campos A, Chalkley RJ, Kraus H-J, Albar JP, Martinez-Bartolomé S, Apweiler R, Omenn GS, Martens L, Jones AR, Hermjakob H. 2014. ProteomeXchange provides globally coordinated proteomics data submission and dissemination. Nat Biotechnol 32:223–226. doi:10.1038/nbt.2839.

