## Supplemental Text S1 for "CP204L Is a Multifunctional Protein of African Swine Fever Virus That Interacts with The VPS39 Subunit of HOPS Complex and Promotes Lysosome Clustering"

### Supplemental Materials and Methods

#### Molecular cloning

##### CP204L-GFP plasmid

For the generation of CP204L-GFP plasmid, the codon-adapted viral frame (ORF) CP204L of ASFV Georgia 2007/1 (Chapman et al. 2011) GenBank accession # FR682468) was amplified from plasmid pUC-BaKJCAG-CP204Lsyn (Hübner et al. 2018) by PCR with primers pCAG-F3 and ASFVp30CDS-R (Table 1) using KOD Xtreme$^{TM}$ Hot Start DNA Polymerase (Sigma/Merck). After digestion of the PCR product with *Bam*HI and *Eco*RI the isolated 617 bp fragment was inserted into the correspondingly digested and dephosphorylated reporter gene expression vector pGFP-N1 (Clontech, GenBank accession # U55762). In the resulting plasmid CP204L-GFP, the synthetic CP204L ORF was under the control of the human cytomegalovirus (HCMV) immediate-early promoter/enhancer complex and 3'-terminally fused to the coding sequence of an enhanced green fluorescent protein (GFP).

##### pVPS39-GFP plasmid

For overexpression of VPS39 and GFP fusion proteins with VPS39 in transfected eukaryotic cells, the predicted gene product of *Sus scrofa* (isoform X3, GenBank accession # XP_013848582) was back-translated in line with porcine codon preferences. The custom-made (GeneArt, ThermoFisher Scientific) synthetic ORF was flanked by a 5'-terminal Kozak sequence (CCACC) and restriction sites for convenient recloning (Fig. 1). For fusion of VPS39 to the N-terminus of GFP, a 2674 bp EcoRI/BamHI fragment was inserted into correspondingly digested pGFP-N1. The obtained precursor plasmid was digested with AhdI, treated with Klenow polymerase, and re-ligated. This treatment caused a frameshift immediately upstream of the stop codon of VPS39, leading to in-frame fusion with the downstream GFP ORF in pVPS39porc-GFP.

**Control plasmid (empty vector)**

To obtain a matching control plasmid for transfection experiments, a 741 bp *Bam*HI/*Not*I fragment containing the ORF encoding the enhanced GFP was deleted from pGFP-N1 (Clontech, GenBank accession # U55762), resulting in pΔGFP-N1 after Klenow-treatment and ligation.

**VPS39 CRISPR/Cas9 knockout cells**

For CRISPR/Cas9 knockout of VPS39, the complementary oligonucleotide pairs VPS39porc-gR3F and VPS39porc-gR3R (Table 1) targeting codons 201 to 208 of the VPS39 splice variant X3 ORF, as well as VPS39porc-gR4F and VPS39porc-gR4R (Table 1) targeting VPS39 codons 100 to 107, were hybridized, phosphorylated and cloned into the BpiI-digested and dephosphorylated single guide RNA (sgRNA) and Cas9 nuclease expression vector pX330A-1x4neoR (unpublished results). After verifying the correct sequence insertions, the resulting plasmids pX330-VPS39gR3neoR and pX330-VPS39gR3neoR were used to transfect WSL cells. After selection with 500 µg/ml G418, single resistant cell clones were propagated, and tested for expression of the FLAG-tagged Cas9 nuclease by western blot analyses of cell lysates using the monoclonal ANTI-FLAG® M2 antibody (Sigma/Merck) as described previously (Hübner et al. 2018). Genomic DNA of Cas9-expressing cell clones was prepared (Fuchs and Mettenleiter 1996) and tested for the presence of the desired sgRNA genes by PCR amplification and sequencing using primers X330GGR-F2 and X330GGR-R2 (Table 1), the BigDye™ Terminator v1.1 Cycle Sequencing Kit, and an Applied Biosystems 3500 Genetic Analyzer (ThermoFisher Scientific). Furthermore, genomic DNA of the cell clones was analyzed for CRISPR/Cas9-induced mutations by PCR amplification and sequencing of the targeted VPS39 gene regions using primers VPS39gR3T-136F and VPS39gR3T-950R, or VPS39gR4T-147F and VPS39gR4T-1003R, respectively (Table 1). Sequences were evaluated using the Geneious Prime 2021.0.1 software package (Biomatters, available from https://www.geneious.com).

These studies revealed that the VPS39 gene of the selected WSL cell clone VPS39-KO1 exhibited a single nucleotide (A) insertion inducing a frameshift behind codon 202 (Fig. 1) and resulting in a premature termination codon at position 224. In contrast, cell clone VPS39-KO2 possessed a single nucleotide (A) deletion in VPS39 codon 101 (Fig. 1) leading to premature termination at position 103. Both cell clones were obviously homozygous with respect to these mutations, and not able to express the CP204L binding domain localized between amino acids 271 and 541 of VPS39.

### Mass spectrometry sample preparation and data analysis

#### Immunoprecipitation

A total of 5 x $10^6$ WSL cells (stably expressing GFP, CP204L-GFP, or VPS39-GFP) were mock-infected or infected with cell culture-adapted ASFV Armenia 2008 at a multiplicity of infection (MOI) of 2 PFU/cell. After 24 hours, cells were washed thrice with PBS and lysed on ice in 1 ml cold immunoprecipitation (IP) buffer [(50 mM Tris-HCl, pH 7.4, 150 mM NaCl, 1 mM $MgCl_2$) supplemented with benzonase (25 U/ml, Sigma-Aldrich #E8263), 0.5% Nonidet P40 substitute (NP-40; Sigma-Aldrich #I8896) and cOmplete mini EDTA-free protease inhibitor cocktail (Roche, #04693159001)]. Cells were lysed by sonication 3x30sec, 80% amplitude (Branson Digital Sonifier 450) and incubated on a tube rotator for 30 min at 4°C and another 30 min on a thermomixer at 37°C with constant shaking at 900 rpm. Lysates were cleared by centrifugation at 13,000 x *g* and 4°C for 15 min. 50 µl of each lysate was removed for immunoblotting (whole cell lysate fraction). GFP-trap agarose beads (50 µl, Chromotek) were washed twice in IP buffer and incubated with the remaining lysate for 1 h with constant rotation at 4 °C. Beads were washed twice with 1 ml of IP buffer containing 0.05% NP-40 followed by two washes in detergent-free IP buffer. 10 µl bead slurry was removed for immunoblotting (IP fraction). The remaining 40 µl beads prepared in at least three independent biological replicates for each bait were kept for mass spectrometric analysis.

**Sample preparation for mass spectrometry**

After IP, beads were suspended in 300 µl freshly made UA buffer [8 M urea, 100 mM Tris-HCl pH 8], loaded onto 10 kDa filter units (Sartorius), and centrifuged at 12,000 x *g* at 20°C for 30 min. Filter Aided Sample Preparation (FASP) trypsin digestion was performed as described previously (Wiśniewski et al. 2009). Beads were trypsinized to digest the baits and the interacting proteins in 100 µl of digest buffer [1 M urea, 50 mM Tris-HCl pH 7.5 and 5 µg/mL Trypsin (Promega)]. Digestion was performed overnight at 37°C with shaking. The next day, the flow-through containing peptides was collected, and the samples were inactivated for 10 min at 95°C. Peptides were acidified with formic acid (1% final concentration) and desalted using C18 100 µl tips (Thermo Scientific) according to the manufacturer's instructions, dried by vacuum centrifugation, and reconstituted in 20 µl of 0.1% formic acid prior to mass spectrometry.

**Protein identification by LC-MS/MS**

Digested peptide mixtures were analyzed by LC-MS/MS on a timsTOF Pro (Bruker Daltonik), which was coupled online to a nanoElute nanoflow liquid chromatography system (Bruker Daltonik) via a CaptiveSpray nano-electrospray ion source. Peptides (corresponding to 100 ng) were separated on a reversed-phase C18 column (25 cm × 75 µm i.d., 1.6 µm, IonOpticks) with a binary buffer system of buffer A (water with 0.1% formic acid, *v*/*v*) and buffer B (acetonitrile with 0.1% formic acid, *v*/*v*). Peptides were separated running a gradient of 2–95% mobile phase B over 115 min (2% to 16% solvent B (0-60 min), 15-24% solvent B (60-90 min), 24%-34% solvent B (90-105 min), 35-95% solvent B (105-107 min) and 95% solvent B (107-115 mins) at a constant flow rate of 400 nl/min. The column temperature was controlled at 40 °C. MS analysis of eluting peptides was performed in ddaPASEF mode (1.1sec cycle time) as recommended by the manufacturer.

**Data analysis**

Mass spectrometry raw files were processed with MaxQuant (v.2.0.2.0) (Cox and Mann 2008). Peptide search was performed against an ENSEMBL (Aken et al. 2017) *Sus scrofa* proteome database (v.11.1.2021-11-10) and an NCBI ASFV Georgia (v.FR682468.2) proteome database. The following modifications were included in the search parameters: trypsin digestion with a maximum of 2 missed cleavages, carbamidomethylation of cysteine as fixed modification, protein N-terminal acetylation and oxidation of methionine as variable modifications. Mass error tolerance was set to 20 ppm for the full scan (MS1) and 40 ppm for MS/MS (MS2) spectra. False discovery rate (FDR) on peptide and protein level was set to 0.01, the minimum peptide length to seven amino acids, and the match between runs option was used with a 0.7 min match window and 20 min alignment time. The output files were analyzed using Perseus software (v.2.0.3.0) (Tyanova et al. 2016). Proteins identified only by modified peptides, reverse hits, and contaminants were filtered out. Protein was considered identified if at least 2 unique peptides of this protein were found in 50% of the replicates.

Further filtering of proteins specifically binding to CP204L protein was achieved by comparing CP204L protein pulldowns with the negative GFP control. iBAQ (intensity Based Absolute Quantitation) protein quantitation data were log2-transformed and filtered to contain a minimum of two valid values in at least one experiment. Missing values were imputed on the total matrix with default settings (width: 0.3, downshift 1.8). Potential interactors were considered CP204L-specific if they were identified only in CP204L pulldowns or the log2 fold change (between control and CP204L) was greater than two, and the *q*-value of a two-sided t-test was <0.01.

**Verification of VPS39 knockout in WSL cells**

The absence of VPS39 in knockout cells was confirmed by LC-MS/MS analysis. Lysates of WSL KO and control cells (100 µg) were prepared using the Thermo EasyPep Mini MS Sample Prep Kit (Thermo Scientific) according to the manufacturer's instructions. Dried peptides were reconstituted in 0.1% formic acid to a final concentration of 100 ng/µl. Peptides corresponding to 200 ng protein were measured by LC MS/MS and analyzed as described in the Data analysis section.

**Silver staining and in-gel digest**

The SDS-PAGE (Laemmli 1970) gel was stained by incubating in colloidal Coomassie stain solution (Neuhoff et al. 1988) overnight. Next, gel was fixed in ethanol (50% in $H_2O$), sensitized with 0.02% $Na_2S_2O_3$ and incubated for 20 min with silver nitrate solution (0.2% $AgNO_3$ + 0.075% formalin in $H_2O$) (Blum et al. 1987). The gel was developed by incubation in a solution of 3% $Na_2CO_3$, 0.0004% $Na_2S_2O_3$ + 0.05% formalin. The reaction was stopped by incubation with 10% acetic acid. Protein bands were excised from gel and placed in a 1.5 ml centrifuge tube to prepare samples for mass spectrometric identification. Next, gel pieces were completely destained in 30 mM $K_3[Fe(CN)_6]$ / 100 mM $Na_2S_2O_3$ : 1/1 and washed with $H_2O$. The gel plugs were washed twice with 50 mM ammonium hydrogen carbonate ($NH_4HCO_3$) and shrunk again with the addition of acetonitrile. The cysteine residues were reduced by incubation in 5 mM dithiothreitol (DTT) at 45°C for 30 min and alkylated by treatment with 22.5 mM iodoacetamide at RT for 20 min. After dehydration with acetonitrile, the proteins were digested in gel overnight with 5 µl of 8 ng/µl of modified porcine trypsin (Promega) in 10 mM $NH_4HCO_3$. The collected supernatant was acidified with formic acid (1% final concentration) and desalted using C18 100 µl tips (Thermo Scientific) according to the manufacturer's instructions, dried by vacuum centrifugation, and reconstituted in 20 µl of 0.1% formic acid prior to mass spectrometry.

**Table 1:** Synthetic oligonucleotide primers

| Name | DNA sequence |
|---|---|
| pCAG-F3 | 5'-GCTAACCATGTTCATGCCTTC-3' |
| ASFVp30CDS-R | 5'-ACA<u>GGATCC</u>GCGATGTACGTCAGGTAGAAGC-3' |

| | |
|---|---|
| VPS39porc-gR3F | 5'-*CACC*GCTCCAGCTGTTTTCCTGTT-3' |
| VPS39porc-gR3R | 5'-*AAAC*AACAGGAAAACAGCTGGAGC-3' |
| VPS39porc-gR4F | 5'-*CACC*GTTGAAATGTCAGTAGGTCG-3' |
| VPS39porc-gR4R | 5'-*AAAC*CGACCTACTGACATTTCAAC-3' |
| X330GRR-F2 | 5'-ATGCTTACCGTAACTTGAAAG-3' |
| X330GRR-R2 | 5'-ATTTGTCTGCAGAATTGGCG-3' |
| VPS39gR3T-136F | 5'-TAGGAATGGGATTGTTGGG-3' |
| VPS39gR3T-950R | 5'-AATGTTGCATCCTTCACCC-3' |
| VPS39gR4T-147F | 5'-ATTGTGCTTGCTTTGTTGG-3' |
| VPS39gR4T-1003R | 5'-GTTTCAACTTTGGCTCACC-3' |

Primers used for generation and characterization of ASFV CP204L expression constructs and of VPS39 knockout swine cells. Relevant restriction sites are underlined, and sequence overhangs for cloning into *Bpi*I-digested pX330A-1x4neoR are printed in Italics.

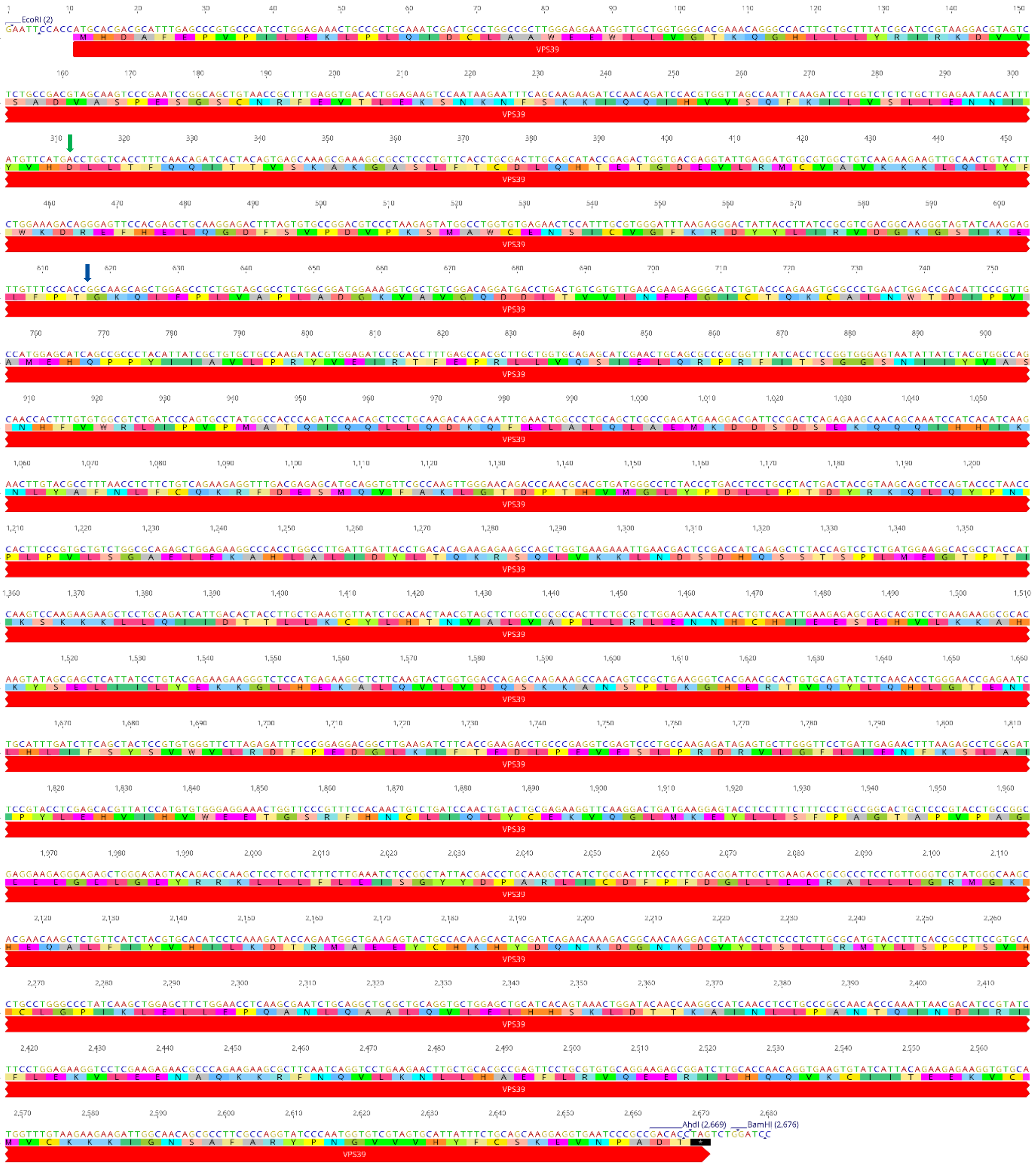

**Fig. 1:** DNA and amino acid sequence of porcine VPS39

The back translated synthetic DNA sequence encoding porcine VPS39 splice variant X3 (accession # XP_013848582) is shown. Relevant restriction sites are indicated. Amino acid positions of VPS39 knockout mutations in WSL VPS39-KO2 (green arrow) and WSL-KO1 (blue arrow) cells are also shown.

### Publication bibliography

Aken, Bronwen L.; Achuthan, Premanand; Akanni, Wasiu; Amode, M. Ridwan; Bernsdorff, Friederike; Bhai, Jyothish et al. (2017): Ensembl 2017. In *Nucleic Acids Research* 45 (D1), D635-D642. DOI: 10.1093/nar/gkw1104.

Blum, Helmut; Beier, Hildburg; Gross, Hans J. (1987): Improved silver staining of plant proteins, RNA and DNA in polyacrylamide gels. In *Electrophoresis* 8 (2), pp. 93–99. DOI: 10.1002/elps.1150080203.

Chapman, David A. G.; Darby, Alistair C.; Da Silva, Melissa; Upton, Chris; Radford, Alan D.; Dixon, Linda K. (2011): Genomic analysis of highly virulent Georgia 2007/1 isolate of African swine fever virus. In *Emerging infectious diseases* 17 (4), pp. 599–605. DOI: 10.3201/eid1704.101283.

Cox, Jürgen; Mann, Matthias (2008): MaxQuant enables high peptide identification rates, individualized p.p.b.-range mass accuracies and proteome-wide protein quantification. In *Nat Biotechnol* 26 (12), pp. 1367–1372. DOI: 10.1038/nbt.1511.

Fuchs, W.; Mettenleiter, T. C. (1996): DNA sequence and transcriptional analysis of the UL1 to UL5 gene cluster of infectious laryngotracheitis virus. In *The Journal of general virology* 77 (Pt 9), pp. 2221–2229. DOI: 10.1099/0022-1317-77-9-2221.

Hübner, Alexandra; Keil, Günther M.; Kabuuka, Tonny; Mettenleiter, Thomas C.; Fuchs, Walter (2018): Efficient transgene insertion in a pseudorabies virus vector by CRISPR/Cas9 and marker rescue-enforced recombination. In *Journal of virological methods* 262, pp. 38–47. DOI: 10.1016/j.jviromet.2018.09.009.

Laemmli, U. K. (1970): Cleavage of structural proteins during the assembly of the head of bacteriophage T4. In *Nature* 227 (5259), pp. 680–685. DOI: 10.1038/227680a0.

Neuhoff, V.; Arold, N.; Taube, D.; Ehrhardt, W. (1988): Improved staining of proteins in polyacrylamide gels including isoelectric focusing gels with clear background at nanogram sensitivity using Coomassie Brilliant Blue G-250 and R-250. In *Electrophoresis* 9 (6), pp. 255–262. DOI: 10.1002/elps.1150090603.

Tyanova, Stefka; Temu, Tikira; Sinitcyn, Pavel; Carlson, Arthur; Hein, Marco Y.; Geiger, Tamar et al. (2016): The Perseus computational platform for comprehensive analysis of (prote)omics data. In *Nat Methods* 13 (9), pp. 731–740. DOI: 10.1038/nmeth.3901.

Wiśniewski, Jacek R.; Zougman, Alexandre; Nagaraj, Nagarjuna; Mann, Matthias (2009): Universal sample preparation method for proteome analysis. In *Nat Methods* 6 (5), pp. 359–362. DOI: 10.1038/nmeth.1322.
